# Benchmarking computational methods to identify spatially variable genes and peaks

**DOI:** 10.1101/2023.12.02.569717

**Authors:** Zhijian Li, Zain M.Patel, Dongyuan Song, Guanao Yan, Jingyi Jessica Li, Luca Pinello

## Abstract

Spatially resolved transcriptomics offers unprecedented insight by enabling the profiling of gene expression within the intact spatial context of cells, effectively adding a new and essential dimension to data interpretation. To efficiently detect spatial structure of interest, an essential step in analyzing such data involves identifying spatially variable genes. Despite researchers having developed several computational methods to accomplish this task, the lack of a comprehensive benchmark evaluating their performance remains a considerable gap in the field. Here, we present a systematic evaluation of 14 methods using 60 simulated datasets generated by four different simulation strategies, 12 real-world transcriptomics, and three spatial ATAC-seq datasets. We find that spatialDE2 consistently outperforms the other benchmarked methods, and Moran’s I achieves competitive performance in different experimental settings. Moreover, our results reveal that more specialized algorithms are needed to identify spatially variable peaks.

## Introduction

Recent years have witnessed significant progress in spatially-resolved transcriptome profiling techniques that simultaneously characterize cellular gene expression and their physical position, generating spatial transcriptomic (ST) data. The application of these techniques has dramatically advanced our understanding of disease and developmental biology, for example, tumor-microenvironment interactions^1^, tissue remodeling following myocardial infarction^2^, and mouse organogenesis^3^, among others.

Spatial transcriptome profiling methods are broadly categorized into two groups, i.e., next-generation sequencing (NGS)-based (including 10x Visium^4^; Slide-seq^5,6^; HDST^7^; STARmap^8^) and imaging-based (including seqFISH^9^ and MERFISH^10^) (**Fig. 1a**). They vary in terms of the number of genes and spatial resolution. Specifically, NGS-based assays usually provide genome-wide gene expression through spots profiling multiple cells, thus precluding the possibility of delineating expression at the single-cell level. At the same time, the imaging-based methods can generate sub-cellular resolution data but can only detect a subset of genes (30-300). Due to these differences in the number of genes and spatial resolution, distinct computational methods and algorithms are required for the downstream analysis of each data type. In the case of NGS-based profiles, an important task involves associating cell types with spatial locations through cell-type deconvolution, often leveraging paired single-cell RNA-seq data to compensate for the low spatial resolution^11–13^. On the other hand, for imaging-based profiles, the initial step involves performing cell segmentation to accurately delineate the boundaries of individual cells^14^.

**Fig. 1.**
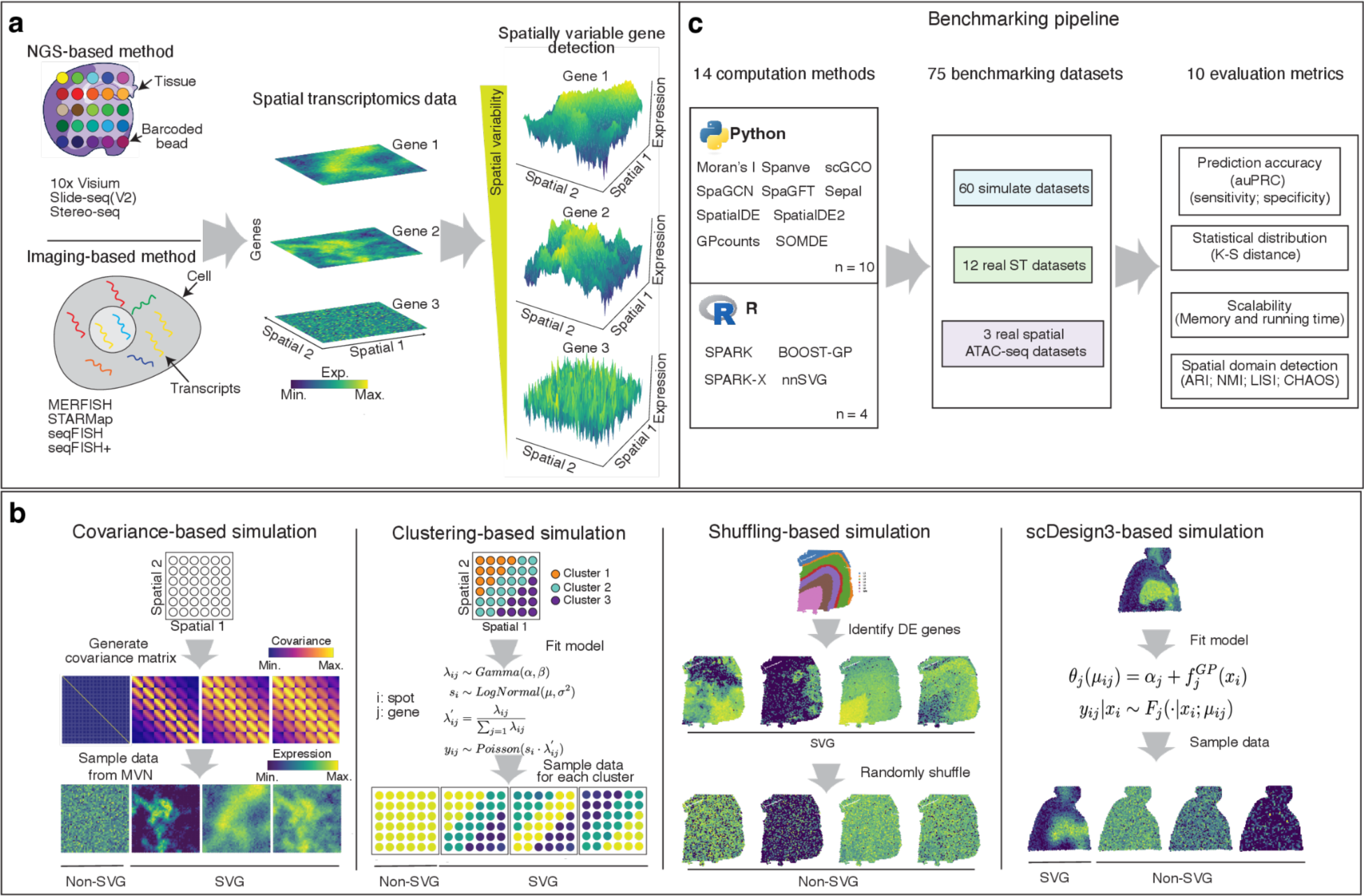
Overview of spatial transcriptome profiling protocols, benchmarking datasets with simulation designs, and benchmarking workflow. a, Left: a schematic showing the NGS-based and imaging-based technologies for profiling spatially resolved transcriptomes. Middle: Visualization of gene expression with various patterns in spatial space. Colors refer to the expression levels of genes. Right: 3D plots showing the expression of the genes with different spatial patterns. A gene with a highly spatially correlated expression pattern is defined as a spatially variable gene (SVG; shown on the top), otherwise as a non-SVG (shown on the bottom). The x-axis and y-axis represent spatial coordinates, and the z-axis represents the expression of that gene. b, Schematics showing four approaches to simulate spatial transcriptomics datasets with ground truths. In the covariance-based simulation, we sampled data from a multivariate normal (MVN) with different covariance matrices for SVGs and non-SVGs. In the clustering-based simulation, we generated SVGs as differentially expressed genes for pre-defined spatial clusters. In the shuffling-based simulation, we first identified cluster-specific DE genes as SVGs and then generated the non-SVGs through data shuffling. In scDesign3-based simulation, we modeled a gene’s expression as a function of spatial locations via Gaussian Process regression. c, Benchmarking workflow. We compared 14 computational methods on 60 simulated, 12 spatial transcriptomics, and three spatial ATAC-seq datasets. The evaluation metrics include prediction accuracy (measured by auPRC, sensitivity, and specificity), statistical distribution similarity (measured by K-S distance), scalability (measured by memory and running time), and spatial domain detection accuracy (measured by ARI, NMI, LISI, and CHAOS). K-S, Kolmogorov-Smirnov.

One common task for all ST profiles, regardless of the employed protocols, is to identify genes that exhibit spatial patterns^15^ (**Fig. 1a**). These genes, defined as spatially variable genes (SVGs), contain additional information about the spatial structure of the tissues of interest, compared to highly variable genes (HVGs). Examples of SVGs include genes involved in developmental gradients^16^, cell signaling pathways^17^, and tumor micro-environment interface^1^. Additionally, SVGs may be critical to downstream tasks such as detecting spatial domains^18^ and inferring spatially aware gene regulatory networks (GRNs)^19^. To detect SVGs, researchers have developed various computational methods by incorporating the spatial context into the analysis. As the number of methods keeps increasing, it becomes difficult for users to choose the best approaches effectively. Previous benchmarking studies have typically compared no more than seven computational methods^20–22^, significantly fewer than the currently available methods (n > 14). Furthermore, since obtaining ground truth from real-world ST profiles is not feasible, these studies have relied on simulation data to evaluate the accuracy of each method in detecting SVGs. However, the simulation data were generated either only using the predefined spatial clusters^20,22^ or with a very limited number of spatial patterns (e.g., *spots* where the expression forms round contours and *linear* where the expression forms rectangular shapes)^21^. Consequently, the limitations of the simulation strategies may introduce inflating performance metrics compared to realistic settings. Therefore, there is a clear need for a comprehensive benchmarking study incorporating more methods and employing enhanced simulation strategies to capture biologically plausible patterns of interest. Such a study would provide a more robust and unbiased evaluation of the available methods for detecting SVGs in ST profiles, enabling researchers to make informed decisions when selecting the most appropriate computational methods for their analyses.

In this work, we comprehensively evaluated 14 methods (see **Table 1**) for identifying SVGs (the selection of the 14 methods is discussed in the Discussion). We created multiple benchmarking datasets (n = 60) with verified ground truths and compared the methods in terms of prediction accuracy, sensitivity, specificity, statistical calibration, and scalability. We also investigated the impact of identified SVGs on spatial domain detection. Finally, we explored the applicability of the methods to other spatial modalities, specifically examining their effectiveness on spatial ATAC-seq data. Our benchmark results indicate that *SpatialDE2*^23^ generally outperformed the other tested methods. Furthermore, *Moran’s I*^24^, despite its simplicity, consistently exhibited performance either comparable to or superior to most methods in our benchmark evaluations. Our results provide a detailed comparison of SVG detection methods and serve as a reference for both users and method developers.

**Table 1.**
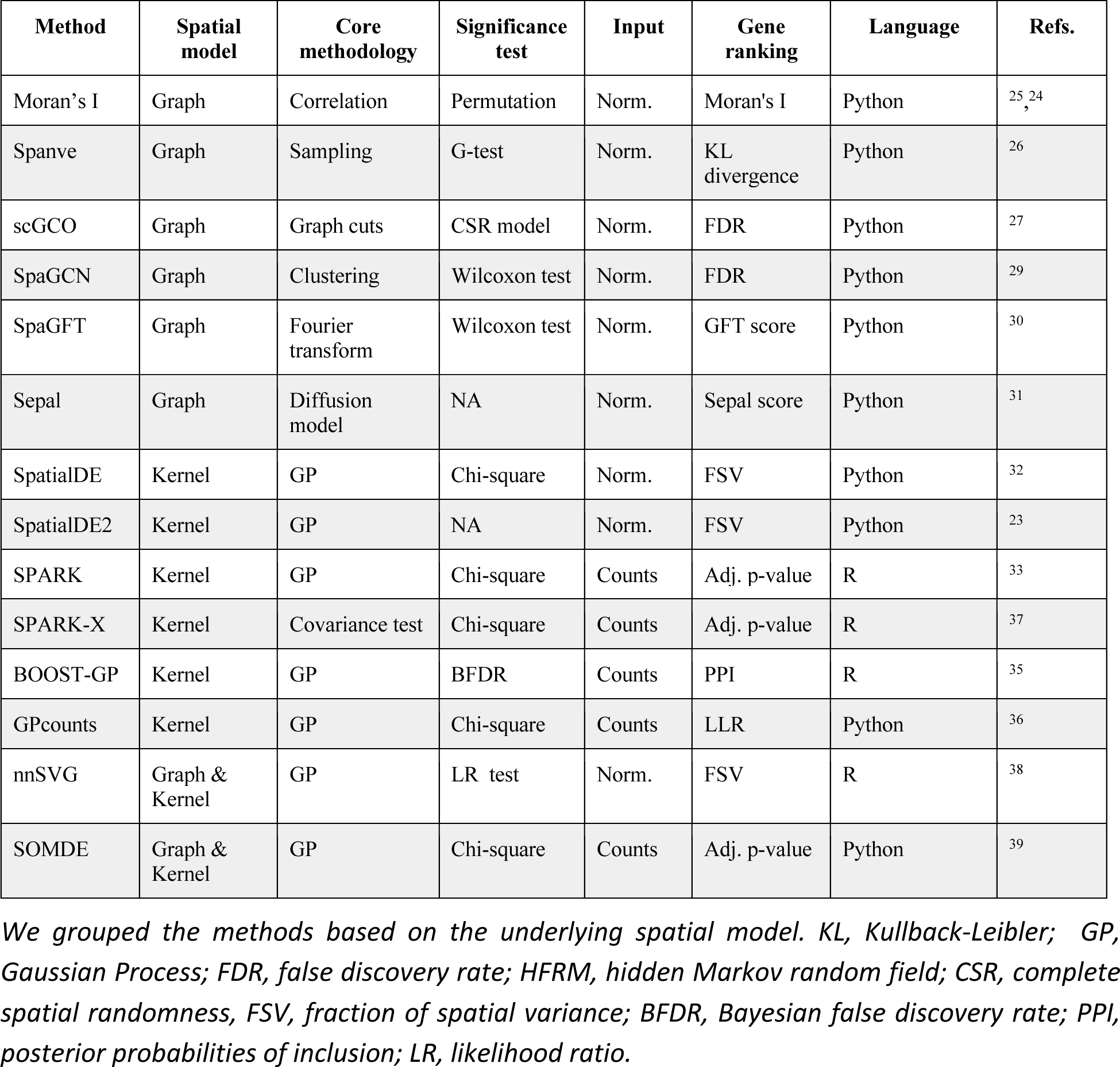
Overview of computational methods for identification of spatially variable genes.

| Method | Spatial model | Core methodology | Significance test | Input | Gene ranking | Language | Refs. |
| --- | --- | --- | --- | --- | --- | --- | --- |
| Moran's I | Graph | Correlation | Permutation | Norm. | Moran's I | Python | <sup>25, 24</sup> |
| Spanve | Graph | Sampling | G-test | Norm. | KL divergence | Python | <sup>26</sup> |
| scGCO | Graph | Graph cuts | CSR model | Norm. | FDR | Python | <sup>27</sup> |
| SpaGCN | Graph | Clustering | Wilcoxon test | Norm. | FDR | Python | <sup>29</sup> |
| SpaGFT | Graph | Fourier transform | Wilcoxon test | Norm. | GFT score | Python | <sup>30</sup> |
| Sepal | Graph | Diffusion model | NA | Norm. | Sepal score | Python | <sup>31</sup> |
| SpatialDE | Kernel | GP | Chi-square | Norm. | FSV | Python | <sup>32</sup> |
| SpatialDE2 | Kernel | GP | NA | Norm. | FSV | Python | <sup>23</sup> |
| SPARK | Kernel | GP | Chi-square | Counts | Adj. p-value | R | <sup>33</sup> |
| SPARK-X | Kernel | Covariance test | Chi-square | Counts | Adj. p-value | R | <sup>37</sup> |
| BOOST-GP | Kernel | GP | BFDR | Counts | PPI | R | <sup>35</sup> |
| GPcounts | Kernel | GP | Chi-square | Counts | LLR | Python | <sup>36</sup> |
| nnSVG | Graph & Kernel | GP | LR test | Norm. | FSV | R | <sup>38</sup> |
| SOMDE | Graph & Kernel | GP | Chi-square | Counts | Adj. p-value | Python | <sup>39</sup> |

We grouped the methods based on the underlying spatial model. KL, Kullback-Leibler; GP, Gaussian Process; FDR, false discovery rate; HFRM, hidden Markov random field; CSR, complete spatial randomness, FSV, fraction of spatial variance; BFDR, Bayesian false discovery rate; PPI, posterior probabilities of inclusion; LR, likelihood ratio.

## RESULTS

### Overview of computational methods for detection of spatially variable genes

In contrast to the identification of HVGs solely from genes expression levels (i.e., mRNA molecular abundance) in single-cell RNA sequencing (scRNA-seq) data, detecting SVGs requires the additional consideration of spatial information at the cellular or subcellular level. A common and straightforward approach is to build a k-nearest-neighbor (KNN) graph where each node represents a spatial spot, and the edges between nodes represent the spatial proximity of spots. SVGs are identified by combining this spatial neighbor graph with gene expression profiles. For instance, *Moran’s I* estimates the correlation coefficient of the expression between a spot and its neighbors^24,25^. Similarly, *Spanve* quantifies the divergence in gene expression distributions between randomly and spatially sampled locations using Kullback-Leibler (KL) divergence^26^. A higher correlation or distribution divergence indicates that the gene is more likely to have a non-random spatial pattern. Moreover, *scGCO* utilizes a hidden Markov random field (HMRF) to capture the spatial dependence of each gene’s expression levels and uses a graph cuts algorithm to identify the SVGs^27^. *SpaGCN* first builds a graph by integrating gene expression, spatial location, and histology information (when available) and then clusters the spots using a graph convolutional network (GCN)^28^; then SVGs are identified by differential expression (DE) analysis on the obtained clusters^29^. *SpaGFT* constructs a KNN graph of spots based on their spatial proximity and then transforms each gene’s expression to the frequency domain; genes with low-frequency signals tend to have less random spatial patterns^30^. *Sepal* uses a diffusion model to assess the degree of randomness of each gene’s spatial expression pattern and ranks the genes accordingly^31^.

Another strategy to incorporate spatial information involves utilizing a kernel function that takes spatial distance as input and computes a covariance matrix to capture the spatial dependency of gene expression across locations. This covariance matrix represents a prior of the underlying spatial pattern. One of the pioneer methods is *SpatialDE*^32^, which models the normalized expression data using non-parametric Gaussian Process (GP) regression and tests the significance of the spatial covariance matrix for each gene by comparing the fitted models with and without the spatial covariance matrix. *SpatialDE2*^23^ further extends this framework by providing technical innovations and computational speedups. *SPARK*^33^ proposes another extension by modeling the raw counts with a generalized linear model based on the over-dispersed Poisson distribution. It provides a more robust statistical approach (Cauchy combination rule^34^) to assess the significance of the identified SVGs. In contrast, *BOOST-GP* uses a zero-inflated negative binomial (ZINB) distribution to model the read counts and infers the model parameters via a Markov Chain Monte Carlo (MCMC) algorithm^35^. Similarly, *GPcounts*^36^ models the counts with a negative binomial (NB) distribution and estimates the model parameters using variational Bayesian inference to improve computational efficiency. Notably, *SPARK-X* stands as an exception by directly comparing the expression covariance matrix and the spatial distance covariance matrix, yielding substantial computational efficiency gains^37^.

In addition, two hybrid methods, namely *nnSVG* and *SOMDE*, have been developed to integrate graph and kernel approaches to capture the spatial dependence between spatial spots. The *nnSVG* method utilizes a hierarchical nearest-neighbor GP to model the large-scale spatial data^38^, providing computational efficiency gains over the standard Gaussian process used in SpatialDE. On the other hand, *SOMDE* employs a self-organizing map (SOM) to cluster neighboring cells into nodes and subsequently fits node-level spatial gene expression using a Gaussian process to identify SVGs^39^. Both methods reduce the computational complexity of kernel approaches by leveraging a spatial graph, which significantly improves their scalability. We summarized the key features of the 14 methods in **Table 1**.

### Benchmarking datasets and pipeline

In this study, the primary challenge we faced while benchmarking the 14 methods for detecting SVGs was the lack of established datasets with verified true labels (i.e., true SVGs and non-SVGs) in real-world scenarios. Hence, we focused on simulated data, an approach grounded in precedent studies^26,27,33,37,38^. Addressing the limitation of previous simulations that predominantly utilized pre-defined spatial clusters — a strategy failing to mirror the rich diversity of spatial patterns —, we formulated three innovative strategies and employed an recent simulation framework to foster a more representative simulation dataset: covariance-based, clustering-based, shuffling-based, and scDesign3-based simulation, illustrated in **Fig. 1b**.

In the covariance-based simulations, we sampled gene expression data from a multivariate normal (MVN) distribution where the covariance matrix was pre-defined based on the spatial coordinates (see Methods). To generate SVGs with various spatial patterns, we employed multiple Gaussian kernels with diverse length scales to define the covariance matrix. We also controlled the noise levels and covariance amplitudes to introduce varying degrees of complexity (**Supplementary Fig. 1**). For non-SVGs, we simply used the identity matrix as the covariance matrix. In the clustering-based simulations, we first fitted a Gamma-Poisson mixture model on real-world spatial transcriptomics profiles from breast tumors with annotated spatial clusters^40^. We then generated synthetic data for each cluster by manipulating the log-fold change for each gene to simulate different gene expression levels and to assess the sensitivity of each SVG detection method (**Supplementary Fig. 2**). In the shuffling-based simulation, we downloaded a spatial transcriptomics dataset generated from the human dorsolateral prefrontal cortex (DLPFC)^41^ with distinct and well-annotated spatial clusters, and we obtained “true labels” based on differential expression analysis and data shuffling. Briefly, we considered the cluster-specific markers as true SVGs and randomly shuffled the spots to remove spatial correlation, creating non-spatially variable expressions (**Supplementary Fig. 3a-c)**. Finally, we used scDesign3^42^, a recent simulation framework for generating realistic spatial transcriptomics datasets with pre-specified true SVGs (**Supplementary Fig. 4).** Details of the simulated datasets were provided in **Supplementary Table 1.**

Using the simulated datasets as described above, we benchmarked 14 SVG detection methods to identify spatially variable genes, covering most of the currently available methods for this task as detailed in **Table 1** (for the method selection, see Discussion). We evaluated their prediction performance based on the area under the Precision-Recall curve (AUPRC), a widely accepted metric for assessing classification accuracy. Additionally, we compared the sensitivity, specificity, and statistical calibration of the methods. Using simulated data, we also investigated the memory requirements and time scalability of the methods in relation to the number of spatial spots. Importantly, considering potential downstream applications, we evaluated the impact of the detected SVGs on spatial domain detection and measured the performance of this task against the true labels using commonly used metrics such as the Adjusted Rand Index (ARI) and Normalized Mutual Information (NMI). Finally, we explored the possibilities of applying these methods, which were developed for spatial transcriptomics data, to spatial ATAC-seq data for detecting spatially variable peaks (SVPs). The results were evaluated based on clustering analysis using the local inverse Simpson’s index (LISI) and the spatial chaos score (CHAOS) metrics^18^. The overall benchmarking workflow is presented in **Fig. 1c**.

### Benchmarking prediction performance of the methods

We reasoned that identifying SVGs can be considered as a binary classification problem where the task is distinguishing SVGs from non-SVGs based on the statistical significance of a calculated score that should capture the degree of spatially variable pattern. Currently available methods typically provide different scores to rank the genes. For example, *SpatialDE* and *SpatialDE2* use the fraction of spatial variance (FSV) estimated by the GP regression model, while *SpaGFT* defined a GFT score as the sum of the low-frequency Fourier coefficients. We first assessed if these scores could correctly separate SVGs from non-SVGs. To this end, we applied the 14 algorithms to 50 simulated datasets generated using different strategies and evaluated the results using auPRC. We observed that the methods exhibited various accuracies across the benchmarking datasets (**Supplementary Fig. 5a-b**). Specifically, for the covariance-, clustering-, and scDesign3-based simulation datasets, most algorithms achieved a high auPRC at a modest noise level, and their performance declined as the noise level increased. On the other hand, for shuffling-based simulation data, we found instead that *SPARK-X*, *SpatialDE2*, *SpaGFT*, and *Moran’s I* showed consistently higher auPRC than alternative methods.

To compare the performance, we ranked the methods based on auPRC for each experimental setting and visualized the overall results within each dataset and across all the datasets (**Fig. 2a; Supplementary Fig. 5c**). Remarkably, we found that *SpatialDE2* outperformed all other methods on two datasets (i.e., covariance- and scDesign3-based simulations), performed the second-best on the shuffling-based simulation data, and performed the third-best on clustering-based simulation data, demonstrating its robust performance. Interestingly, our evaluation revealed that *Moran’s I* statistic, which solely relies on auto-correlation between spots and their neighbors, showed the second-best performance despite its relative simplicity compared to other methods (**Supplementary Fig. 5c).** Moreover, *SPARK* and *SPARK-X* displayed similar performance, likely because they used the same kernel functions to capture spatial dependency.

**Fig. 2.**
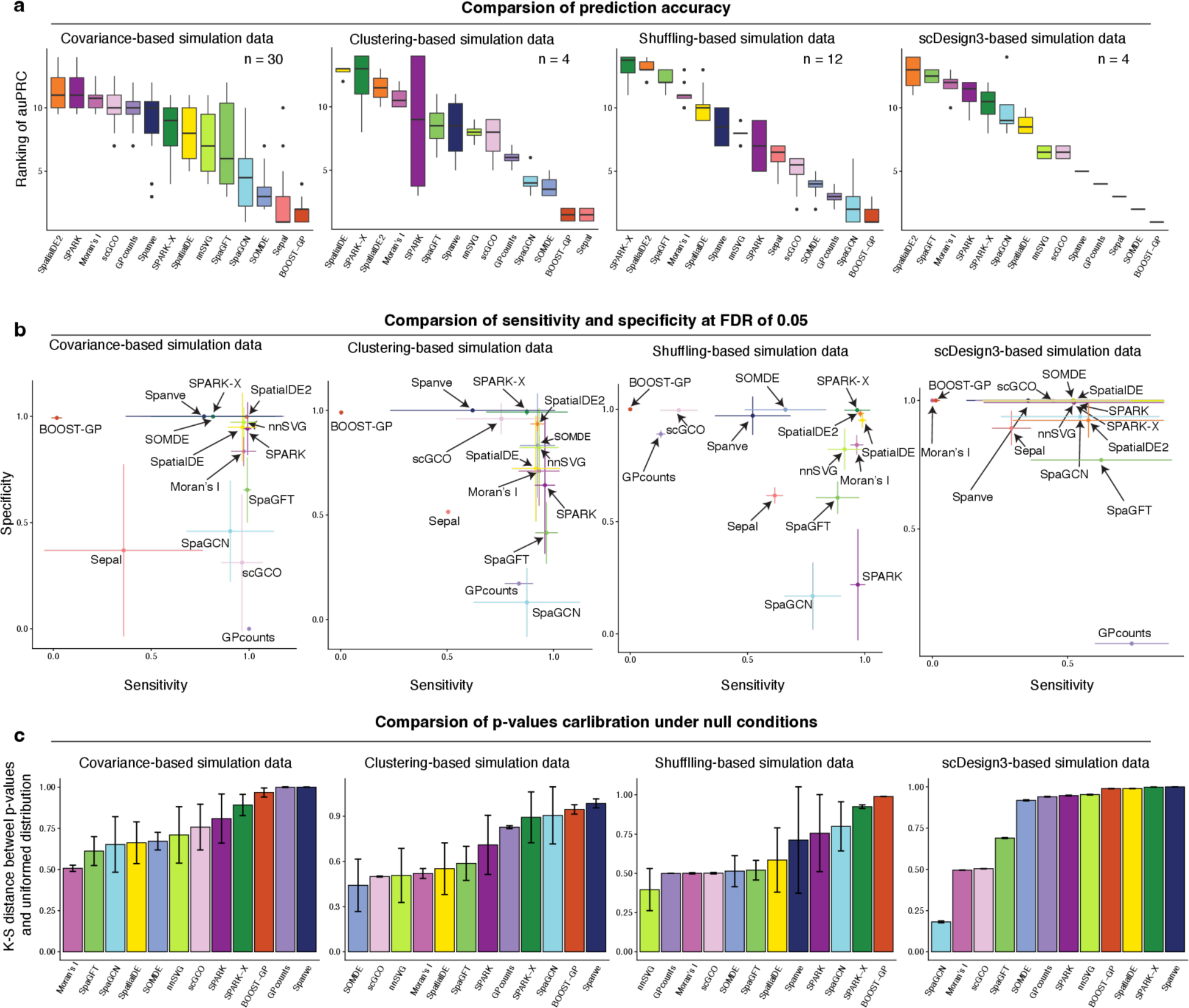
Comparison of the methods using simulated datasets. **a,** Box plot comparing the prediction performance of the methods for covariance-based (n=30), clustering-based (n=4), shuffling-based (n=12), and scDesign3-based (n=4) simulation datasets. The y-axis represents the rank of the method based on auPRC. A higher rank denotes a higher auPRC. **b,** Evaluation of sensitivity and specificity of each method for a false discovery rate (FDR) of 0.05. Each dot represents an average true positive rate and a true negative rate. The error bar represents the standard deviation of the corresponding values. **c,** Bar plot comparing the statistical calibration evaluated by K-S distance between the distribution of empirical p-values and uniformed distribution for the null hypothesis. A lower K-S distance represents a more calibrated model. K-S: Kolmogorov–Smirnov.

### Sensitivity, specificity, and statistical distribution under null conditions

Many of the benchmarked methods also calculate statistical significance, enabling users to identify the most relevant SVGs *ad hoc*. However, because they utilize distinct statistical tests based on different null hypotheses, it is unclear how sensitive and specific the results are. To address this, we subsequently analyzed the methods’ sensitivity (true positive rate) and specificity (true negative rate) at a false discovery rate (FDR) of 0.05. Of note, *SpatialDE2* and *Sepal* were excluded from this analysis because they do not provide statistical significance results.

In terms of sensitivity, we found that most methods achieved high values on the datasets with low noise. However, the performance decreased when the noise level increased, a trend that mirrored our findings in the accuracy evaluation (**Supplementary Fig. 5a**). Intriguingly, our analysis showed that no single method consistently outperformed the rest in both sensitivity and specificity across all benchmark datasets (**Fig. 2b**). For example, *SPARK* and *SpaGFT* displayed high sensitivity but low specificity. In contrast, *Spanve* and *SOMDE* showed high specificity but low sensitivity (**Supplementary Fig. 5-6**). These findings suggest that more sophisticated statistical approaches are needed to control both false positives and false negatives. Nevertheless, we found that *SpatialDE* exhibited the best balance between sensitivity and specificity.

Additionally, we evaluated the p-value distribution of the methods under null conditions for each dataset. To this end, we measured the Kolmogorov–Smirnov (K-S) distance between the distribution of the computed p-values and the uniform distribution (ranging from 0 to 1). The intuition is that a well-calibrated model should produce uniformly distributed p-values between 0 and 1 under the null condition. Therefore, a smaller distance represents a better-calibrated approach. Our analysis revealed that the methods demonstrated various degrees of calibrations in different datasets (**Fig. 2c; Supplementary Fig. 8a-b**). For instance, *SOMDE* showed the best calibration on the clustering-based simulation dataset but was not well-calibrated on the other three datasets. Next, we aggregated the results across all the benchmarking datasets to compare the methods comprehensively. We found that *Moran’s I* exhibited the best calibration among the selected methods. This can potentially be attributed to that this method used permutation to estimate the background distribution, thereby accurately recapitulating the true negatives (i.e., non-SVGs).

### Benchmarking the scalability of the methods

Subsequently, we evaluated the space and time scalability of the analyzed methods. Given that all methods independently estimate the spatial variability for each gene, the scalability, in theory, is primarily influenced by the number of spatial locations. To benchmark this aspect, we generated ten simulation datasets, each consisting of the same number of genes (n = 100) but varying the number of spots, ranging from 100 to 40000. We applied every method to each of the ten simulation datasets and recorded the memory consumption and running time as performance metrics (**see Methods**).

Our initial examination of memory usage revealed that most methods displayed moderate memory requirements, typically staying below 32 GB, even when confronted with datasets containing 40000 spots (**Fig. 3a**). For example, we observed that *Moran’s I* consumed less than 4GB for all datasets. This favorable outcome suggests that these methods can be executed on modern laptops without encountering memory constraints. Among them, *SOMDE* exhibited the most efficient memory usage across all benchmarking datasets, followed by *Spanve* and *SPARK-X* (**Supplementary Fig. 9a**). In contrast, both *SPARK* and *SpatialDE* exhibited significant increases in memory demand as the number of spots in the dataset increased. For instance, when applied to a dataset with 20000 spots, *SPARK* necessitated approximately 250 GB of memory, while *SpatialDE* consumed roughly 150 GB when dealing with a dataset containing 40000 spots. These observations can be attributed to the fact that both *SPARK* and *SpatialDE* are based on Gaussian Process regression, requiring the estimation of a covariance matrix across all spots. Consequently, this leads to a cubic scaling relationship with the number of spots, resulting in the pronounced memory consumption we observed for these two methods.

**Fig. 3.**
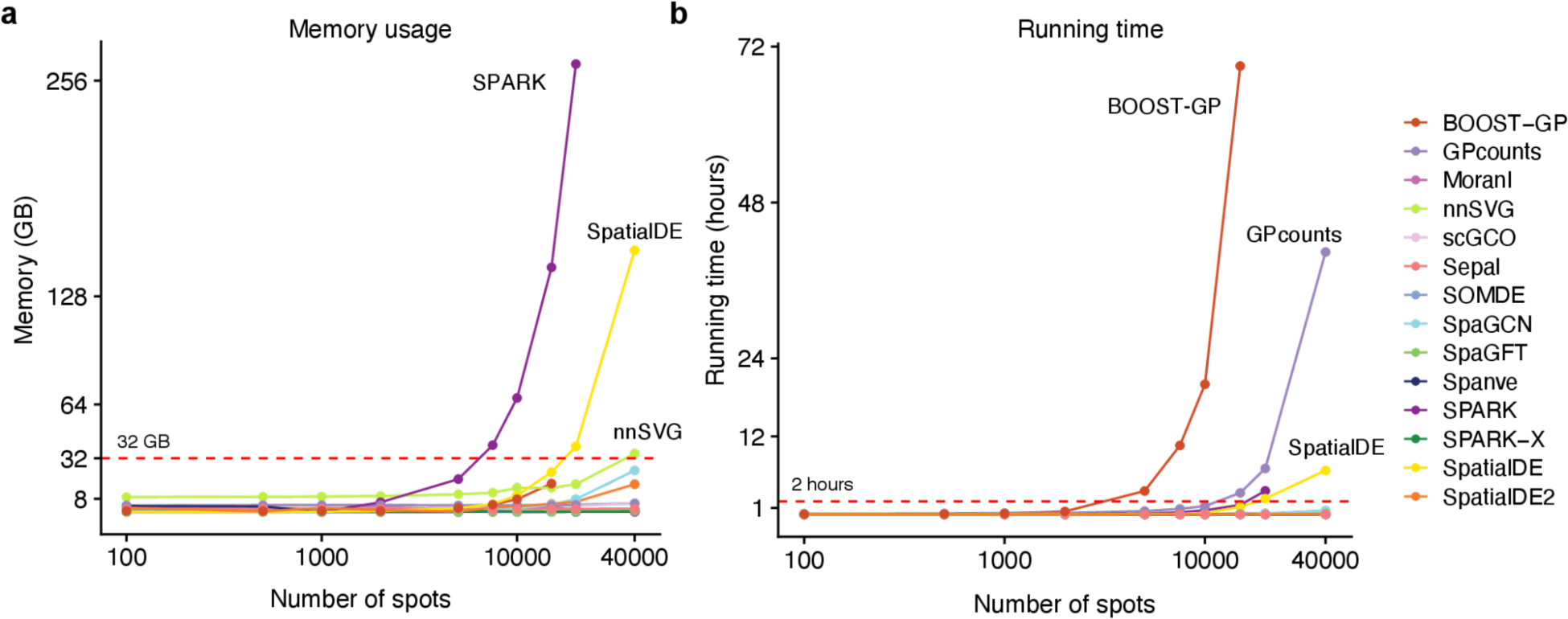
Scalability of the methods. **a,** Line plot showing the memory scalability of the methods. The x-axis represents the number of spots (log10) of the input datasets with 100 genes. The y-axis represents consumed memory (measured as GB) by each method. The red dash line denotes 32 GB. We labeled the top four methods. **b,** Same as **a** for time scalability. The y-axis represents the consumed time (measured as hours) of each method.

Regarding running time, we observed that *SOMDE* achieved the best scalability again, closely followed by *SPARK-X* and *scGCO*. Notably, most methods completed their computations within a reasonable timeframe of about 2 hours (**Fig. 3b; Supplementary Fig. 9b**), making them suitable for practical usage. Both *BOOST-GP* and *GPcounts* exhibited poor scalability with increasing numbers of spots. For instance, *BOOST-GP*’s computational time escalated significantly, requiring three days to process a dataset containing 20000 spots and failing to produce results within five days for a dataset with 40000 spots. Similarly, despite running on a GPU, *GPcounts* still require approximately 45 hours to process the largest datasets. In summary, our analysis revealed that *SOMDE* and *SPARK-X* exhibited the most favorable scalability when handling datasets with an increasing number of spots. In addition, *SpatialDE2* and *Moran’s I* statistics, the top two performers in the evaluation of prediction accuracy, also demonstrated competitive scalability.

### Benchmarking the impact of identified SVGs on spatial domain detection

One of the important applications of spatially resolved transcriptomics is the identification of tissue or region substructures through domain detection analysis. In non-spatially resolved scRNA-seq data, it is a standard practice to utilize HVGs as features for cell clustering^43^. Therefore, we hypothesized that employing SVGs could similarly be beneficial for spatial domain detection. Using the human DLPFC datasets, we first evaluated which method might capture the most informative features for this task. To this end, we ran the methods to identify SVGs for each dataset and observed significant variations in the number of detected SVGs, highlighting discrepancies between the methods (**Supplementary Fig. 10a**). Specifically, *scGCO*, *SOMDE*, *GPcounts*, and *Spanve* tended to yield a low number of SVGs (<1000) across all datasets. In contrast, *SpaGFT*, *Moran’s I*, *SpaGCN*, and *SpatialDE* generated more SVGs. Because *SpatialDE2* and *Sepal* do not perform significance test, we here used the top 2000 genes based on the FSV and *Sepal* scores, respectively (see Methods).

Subsequently, we used the graph-based Leiden algorithm ^44^ (resolution = 1) to cluster the spatial spots based on the detected SVGs as input features for each method and dataset. To establish a baseline for comparison, we also selected the top 2000 HVGs, therefore discarding spatial information in this feature selection procedure. The detection results were evaluated against the annotated spatial domains using two metrics: Adjusted Rand Index (ARI) and Normalized Mutual Information (NMI) (see Methods for details). Remarkably, we observed that all methods, except for *GPcounts* and *Sepal*, exhibited improved accuracy when utilizing SVGs compared to using only HVGs (**Fig. 4a-b; Supplementary Fig. 10b**). This finding underscores the importance and power of incorporating spatial information into this analysis, which can better capture the spatial organization and tissue structures. Among the evaluated methods, *SpatialDE2* demonstrated the highest average ARI (0.31) and NMI (0.44), further confirming its superior performance. Additionally, *Moran’s I* achieved the second-highest average ARI (0.303), closely followed by *SPARK* (0.301) and *SPARK-X* (0.296). Concerning the NMI metric, *SPARK* ranked second-best with an average value of 0.438, followed by *nnSVG* (0.434) and *SPARK-X* (0.43). In conclusion, our analysis highlighted that incorporating SVGs can notably enhance clustering accuracy in spatial transcriptomic analysis. Furthermore, it revealed that *SpatialDE2*, *SPARK*, and *SPARK-X* generally outperformed other methods in this context, showcasing their effectiveness in capturing meaningful spatial patterns and facilitating the discovery of tissue structures in spatial transcriptomics data.

**Fig. 4.**
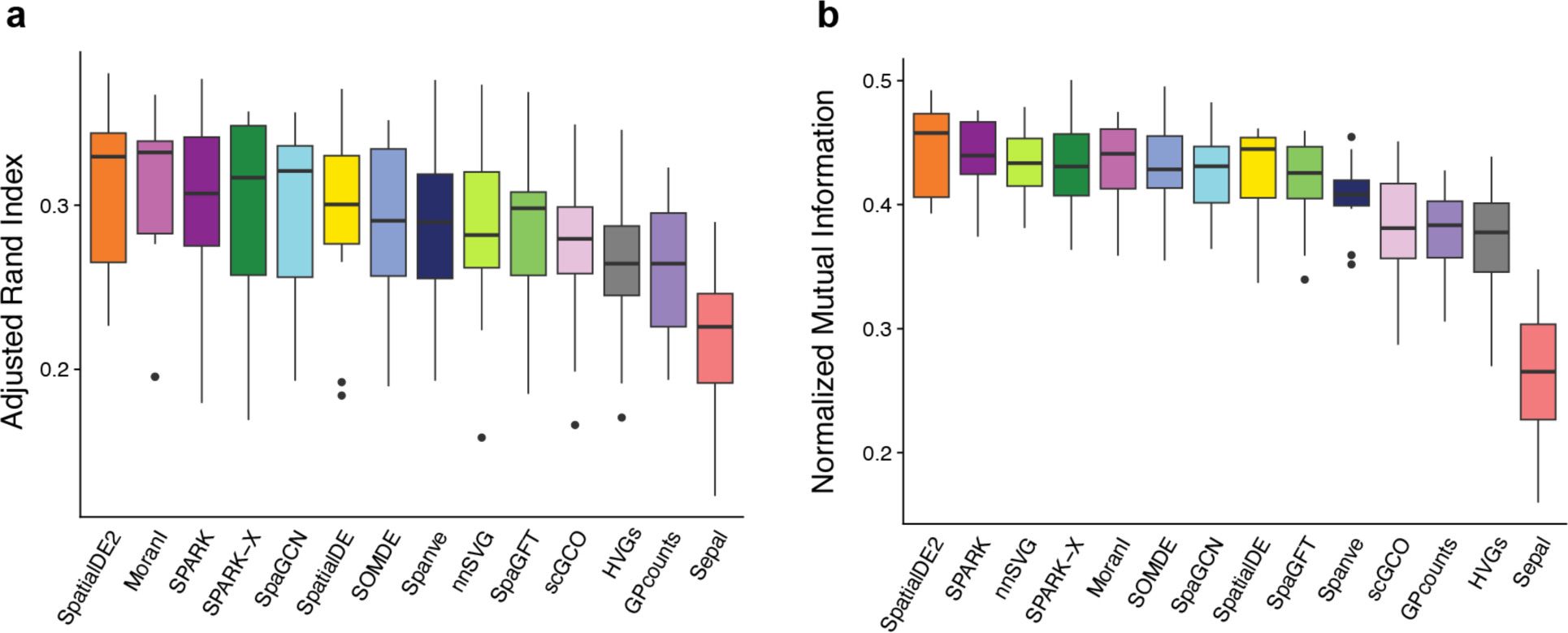
Impact of detected SVGs on spatial domain detection analysis. **a,** Box plots showing the clustering performance as evaluated using ARI. Methods are ranked by the average value. **b,** Same as **a** for NMI.

### Benchmarking the methods with spatial ATAC-seq profiles

Recent technological advances have allowed for profiling spatially-resolved chromatin accessibility^45,46^. However, specific methods for detecting spatially variable open chromatin regions (i.e., spatially variable peaks, abbreviated as SVPs) are currently lacking. In this section, we aimed to investigate the feasibility of applying methods developed for SVG detection to analyze spatial chromatin accessibility profiles. For this, we downloaded spatial ATAC-seq data from mouse gestational development at embryonic days of E12.5^46^ (**Fig. 5a**). Following data processing, we obtained a dataset consisting of 2246 spatial spots and 34460 peaks representing open chromatin regions (see **Methods**). Subsequently, we tried to employ each of the 14 methods to detect SVPs. However, given that these methods were not specifically designed for this task, we encountered several challenges. *BOOST-GP* and *GPcounts* failed to produce results even after 120 hours of running, due to the fact the number of peaks exceeded substantially the number of genes, highlighting the limitation of these two methods in terms of scalability. Additionally, *SPARK* encountered memory issues and did not yield any results. As in the previous section, we wanted to investigate if SVPs recovered from these procedures could boost spatial clustering. Since *SpatialDE2* and *Sepal* do not provide statistical results, we here used the top 20000 peaks. For other methods, we determined the peaks at the FDR of 0.05. Importantly, we observed considerable variation in the number of SVPs detected by different methods (**Fig. 5b**). For example, *nnSVG* and *SOMDE* did not identify any significant peaks, indicating their limitations in capturing spatial variability in this context. In contrast, *SpaGFT* identified almost all the peaks as significant as SVPs (32079 out of 34460)

**Fig. 5.**
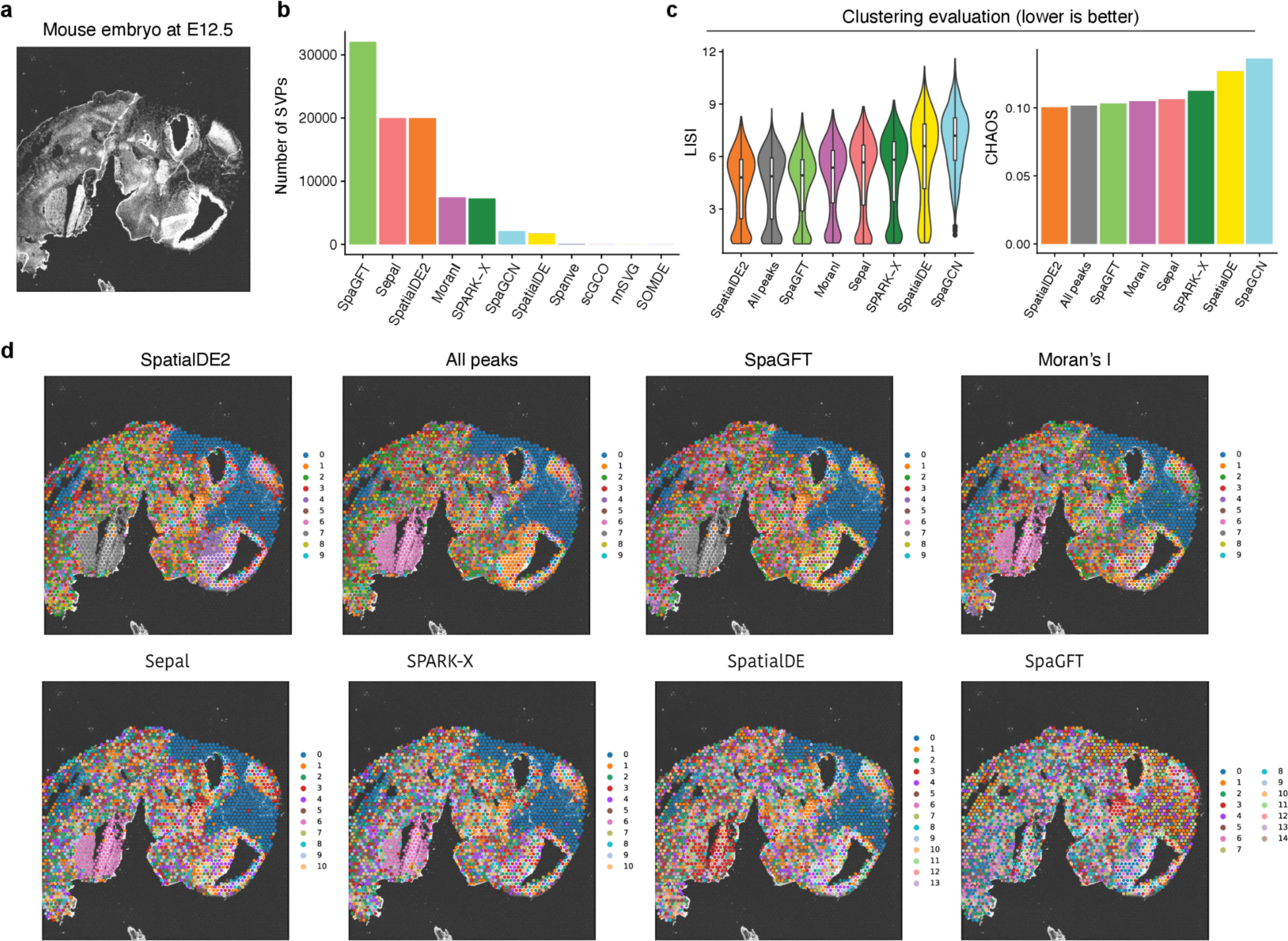
Benchmarking the methods on spatial ATAC-seq data. **a,** Image of a mouse embryo at days of E12.5. **b,** Number of detected spatially variable peaks by each method. **c,** Left: violin plot showing the LISI scores. Methods are sorted by the median values. Right: The bar plot shows the CHAOS score. For both metrics, a lower value represents a better performance. **d,** Visualization of obtained clusters by using spatially variable peaks identified by different methods.

In the subsequent step, we used Leiden-based clustering analysis—utilizing the SVPs to group the spots—to evaluate the quality of SVPs discovered by each method. We excluded *Spanve* and *scGCO* for this analysis as they only detected 39 and 26 SVPs. Because the ground truth is unavailable in this dataset, we measured the spatial locality and continuity of the clusters using two metrics: the local inverse Simpson’s index (LISI) and the spatial chaos score (CHAOS)^18^. The underlying assumption is that a more accurate identification of SVPs would yield more continuous and cohesive clusters^18^. We also included the results generated using all the peaks as a baseline. Interestingly, we observed that *SpatialDE2* outperformed other methods (median LISI = 4.8; CHAOS = 0.1), indicating that it has good potential to identify SVPs (**Fig. 5c-d**). Surprisingly, our analysis revealed that using all peaks yielded the second-best performance (median LISI = 4.87; CHAOS = 0.102). This finding suggests that more specialized methods are required to analyze spatial chromatin accessibility data. Similar results were also observed from the spatial ATAC-seq from embryos at E13.5 and E15.5 (**Supplementary Fig. 11**).

## Discussion

Recently, over a dozen computational methods have been developed to identify spatially variable genes for spatial transcriptomics data. These methods diverge substantially in several aspects, including the assumptions in modeling spatial relationships between cells (graph vs. kernel), the algorithms to estimate spatial variation (e.g., auto-correlation vs. Gaussian Process regression vs. graph cut), the statistical tests to determine significances (e.g., permutation test vs. Wilcoxon test vs. Chi-square test), the choice of input data (raw counts vs. normalized data), and the programming languages (Python vs. R) (**Table 1**). These factors complicate the selection of methods for users, a situation exacerbated by the current absence of systematic benchmarking of the methods’ performance.

In this study, we systematically evaluated the performance of 14 methods for detecting SVGs using simulated and real-world data. Compared to previous works^20,22^, we used four different approaches to generate simulation data. The rationale behind this quadruple simulation strategy is to minimize potential biases and prevent the undue advantages of certain methods on specific types of simulated data. For instance, methods like *SpatialDE*, which models spatial covariance in their algorithmic framework, might overestimate performance when evaluated on covariance-based simulation data. On the other hand, methods such as *SpaGCN* can benefit from clustering-based simulation as this method utilizes clusters to identify SVGs. Moreover, the shuffling-based simulation enables the testing of methods against real-world spatial transcriptomics data. In addition, scDesign3-based simulation can generate *in silico* spatial transcriptomics data, enhancing our simulation with a method capable of explicitly accounting for dependencies between genes. Overall, our benchmark datasets covered a variety of scenarios and represented a useful resource for developing and testing methods in the future.

Our evaluation results revealed that *SpatialDE2* generally outperformed other methods by providing a high average auPRC across various experimental settings, however, this method does not provide any statistical significance for the recovered genes. Interestingly, we found that *Moran’s I* achieved the second-best prediction performance, although it is simply based on auto-correlation between spots and their spatial neighbors, which has been neglected in previous benchmarks^23,26,27,30,32,33,37^. Going forward, it would be prudent to include *Moran’s I* as a baseline in future SVG benchmarking. Additionally, we observed comparable effectiveness between kernel-based and graph-based methods on simulation data and real-world datasets, suggesting their capability to effectively capture similar spatial dependencies. Regarding the sensitivity and specificity, we observed that no single method consistently outperformed the others for both metrics on all benchmarking datasets, indicating that robustly estimating statistical significance remains a difficult problem. Our analysis highlighted the superior p-value calibration of *Moran’s I*, which is attributable to their use of permutation tests that produce well-calibrated statistical significance.

Scalability is another crucial aspect to consider, especially with the emergence of large-scale spatially-resolved profiling methods capable of capturing sub-cellular resolution and accommodating an increasing number of spots, exemplified by Stereo-seq^3^ (> *10*^4^ spots). Our investigations revealed notable distinctions in scalability between graph-based and kernel-based methods, with the former generally outperforming the latter. Among the methods we examined, *SOMDE* stood out as the most efficient in both memory utilization and running time (**Supplementary Fig. 9**). It is important to note that *SOMDE* initially clusters adjacent data points into graph nodes and then employs GP regression to identify SVGs in a node-centric manner. This strategy significantly mitigates the complexities associated with both time and memory. SPARK-X, as a kernel-based method, demonstrated comparable performance to *SOMDE* by directly comparing the expression and spatial distance covariance matrix rather than using GP regression to estimate spatial variation, unlike its predecessor *SPARK*. Moreover, we found that *SpatialDE2* demonstrated reasonable scalability. Given its superior prediction performance, we envision that this method could be the default one to use in practice. Of note, this method provides no statistical significance for its SVG identification results. Therefore, we recommend selecting the top genes based on the fraction of spatial variation (FSV) for downstream analysis. In summary, our findings not only underscore the significance of scalability in the context of SVG detection but also shed light on the relative advantages of different analytical methods when processing large-scale datasets. Future methods should consider scalability alongside prediction performance as advanced spatial profiling techniques produce better quality and larger quantity of data.

In addition, we also demonstrated that the incorporation of SVGs identified by 12 out of the 14 methods led to a notable enhancement in spatial domain detection when applied to real data with annotated clusters, as opposed to relying solely on HVGs. These results imply the significance of capturing spatial information in improving clustering analysis by incorporating more comprehensive information on the architecture of complex tissues and tumors. As novel technologies like Slide-Tag^47^ emerge, enabling the simultaneous acquisition of single-cell measurements and spatial data, we anticipate a surge in the adoption and popularity of SVG identification tools in various downstream analysis tasks.

Finally, we also showed that some methods can be applied to other modalities like Spatial-ATAC seq, facilitating the identification of potential SVPs. We use the term “potential” due to the absence of a ground truth; instead, we leveraged the SVPs for clustering and evaluated the spatial locality and continuity of the obtained clusters. It is essential to note that not all methods were capable of detecting SVPs due to limitations in memory or algorithmic complexity. Therefore, there is a pressing need to develop novel methodologies or modify existing ones to make them applicable to spatial-ATAC seq data. Tools focused on discerning SVPs have the potential to reveal the regulatory elements that govern gene expression profiles within specific spatial sub-regions. This, in turn, can enhance our understanding of the regulatory mechanisms governing SVGs and, consequently, the spatial organization of tissues and tumors. In the future, integrating SVGs and SVPs through novel algorithms holds tremendous potential to facilitate the construction of accurate spatially aware gene regulatory networks.

Although we have covered a large number of available methods (n = 14) in the present study, there are still some methods that are not included. This is because either the repository has not been maintained for a long time, resulting in outdated dependencies that make it difficult to install and execute, for instance, *trendsceek*^48^, or the method was unavailable during the preparation of our manuscript, for example, *BSP*^49^. Another limitation of our work is its exclusive focus on spatial transcriptomics and spatial ATAC-seq, despite the advent of other spatially-resolved omics data, including spatial proteomics. Future directions may also include testing and or adapting SVG detection methods on these modalities. Nonetheless, our benchmarking study provides a detailed evaluation of various SVG detection methods across simulated and real-world datasets of spatial transcriptomics and spatial-ATAC-seq.

## Methods Simulation datasets

We used for different approaches to generate simulated spatial transcriptomics data with ground truth. The details are described below.

### Covariance-based simulation

This simulation is based on a pre-defined covariance matrix. Specificially, given *m* spatial locations where each location is represented by its coordinates *x*, for each gene, we first calculated a covariance matrix *K* ∈ *R^m^*^×*m*^ based on multiple Gaussian kernels:

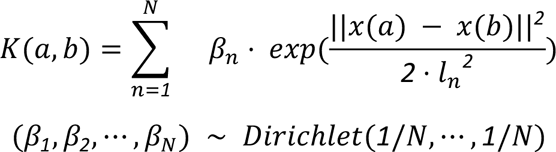

where *x*(*a*) and *x*(*b*) denote two spatial locations, *N* is the number of kernels, β*_n_* is the weight of the *n*th kernel and is sampled from a Dirichlet distribution, *l*_*n*_denotes the length scale. By sampling β for each gene, we obtained different spatial covariance matrices.

We next sampled expression *λ_j_* ∈ *R*^*m*^ for gene *j* across all locations from a multivariate normal distribution (MVN) as follows:

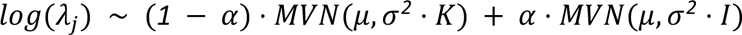

where σ^2^ represents the amplitude of the spatial covariance; *I* is an identity matrix (i.e. with zeros everywhere except on the diagonal); α ∈ [*0*, *1*] denotes the noise level in simulated gene expression. When α = *1*, signals are sampled from an MVN without spatial correlation, thus they are considered non-spatially variable genes. Because some methods can only work on raw counts, we next converted the data to counts as follows:

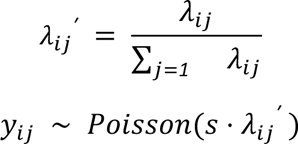

where *s* denotes the library size and is set to 10,000 for all locations.

To evaluate the prediction accuracy, we generated simulation data on a 50 by 50 grid layout (in a total of 2500 spots) by setting *N* = *5* and using *l* ∈ [*1*, *3*.*25*, *5*.*5*, *7*.*75*, *10*] to generate different covariance matrices. Moreover, we used five noise levels α ∈ [*0*, *0*.*2*, *0*.*4*, *0*.*6*, *0*.*8*]and six different amplitudes of the covariance matrix σ ∈ [*0*.*5*, *1*, *1*.*5*, *2*, *2*.*5*, *3*] to generate 30 simulation datasets (**Supplementary Fig. 1**). We generated 100 SVGs and 100 non-SVGs for each dataset using the above process.

To benchmark the scalability of the methods with the number of spatial spots, we generated ten simulation datasets as described above. Each dataset had the same number of genes (n = 100) and a different number of spots (n = 100, 500, 1000, 2000, 5000, 7500, 10000, 15000, 20000, 40000).

### Clustering-based simulation

This simulation is based on spatial data with annotated clusters. To generate a simulation that can recapitulate a real ST dataset, we followed the two-step strategy proposed in SRTsim^50^, i.e., first estimating the parameters required for the simulation from a real ST dataset and then generating a synthetic dataset based on the estimated parameters. Specifically, given a real-world dataset with a count matrix *Y* ∈ *R*^!×*n*^ where *m* is the number of spatial locations, *n* is the number of genes, and *y_ij_* is the expression of gene *j* in spatial location *n*, we modeled the count *y_ij_* using the following process:

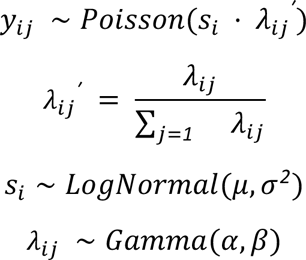

We denoted *s_i_* the total number of reads in location *i* and assumed that it follows a Log-Normal distribution parameterized by μ and σ. We denoted *λ*_*ij*_ the log normalized mean expression of gene *j* sampled from a Gamma distribution parameterized by α and β. *λ*_*ij*_ *^ʹ^* represents the proportion of gene *j* at the location *i*. We estimated the parameters μ, σ, α, and β using the maximum likelihood algorithm based on the count matrix *Y* and log normalized matrix *Y*_*n*()!_ which was generated using functions pp.normalize_total and pp.log1p from the scanpy package^51^.

Once we inferred the parameters, we used them to generate synthetic data by sampling data using the above process. To obtain SVGs, we randomly selected a number of genes for each input spatial cluster and multiplied the sampled mean expression by a differential factor. Since the clusters are spatially associated, these marker genes are considered as SVGs. For non-SVGs, the differential factors were set to one. Using breast tumors as input, we generated simulation data with 100 SVGs and 100 non-SVGs. We varied the differential expression levels from 0.5 to 2, generating four simulation datasets (**Supplementary Fig. 2**).

### Shuffling-based simulation

To test the methods against real-world data, we here created true labels through data shuffling. For this, we downloaded the LIBD human dorsolateral prefrontal cortex (DLPFC) spatial transcriptomics data from http://research.libd.org/spatialLIBD. The data was generated with the 10x Genomics Visium platform and included 12 samples. Each sample was manually annotated as one of the six prefrontal cortex layers (L1-6) and white matter (WM). We filtered the genes by the number of detected spots (>500). Next, we identified marker genes for each cluster with differential expression analysis (t-test, p-value < 0.01, and logFC > 1). These marker genes were considered true positives (i.e., spatially variable genes). Next, we randomly permuted the spots to remove spatial correlation to generate uniformly distributed gene expression profiles. We considered these genes as true negative (i.e., non-spatially variable genes). This resulted in an average number of 549 true labels across all the samples (**Supplementary Fig. 3**).

### scDesign3-based simulation

scDesign3 aims to generate realistic in silico data by first learning interpretable parameters from real data and then generating synthetic data. We installed scDesign3 (v0.99.6) from https://github.com/SONGDONGYUAN1994/scDesign3 and followed the tutorial to generate four datasets with different numbers of true positives ranging from 50 to 200 (**Supplementary Fig. 4**).

### Identify SVGs with computational methods

We described below the details of running the methods to identify SVGs.

#### Moran’s I

*Moran’s I* measures the correlation of gene expression between a spatial location and its neighbors^25^. We computed *Moran’s I* score using Squidpy (v1.2.3)^24^ by following the tutorial: https://squidpy.readthedocs.io/en/stable/auto_examples/graph/compute_moran.html. Spatial neighbors were found using the function spatial_neighbors, and scores were estimated using the function spatial_autocorr. We set parameter n_perms to 100 to obtain the statistical significance and used 0.05 as the threshold for the adjusted p-value to identify significant SVGs. To compute auPRC, we used the *Moran’s I* score to rank genes.

#### Spanve

*Spanve* (Spatial Neighborhood Variably Expressed Genes) is a non-parametric statistical approach for detecting SVGs^26^. Similar to *Moran’s I*, this method uses the difference between a location and its spatial neighbors to estimate the spatial variation. Specifically, for each gene, it computes Kullback-Leibler divergence between space-based and randomly sampled expressions. The significance is calculated by the G-test. We installed *Spanve* (v0.1.0) and ran the method by following the tutorial: https://github.com/zjupgx/Spanve/blob/main/tutorial.ipynb. Genes were ranked by FDR to compute auPRC. We used 0.05 as the threshold for FDR to select significant SVGs.

#### SpaGFT

*SpaGFT* is a hypothesis-free Fourier transform model to identify SVGs^30^. It decomposed the signal from the spatial domain to the frequency domain based on a spatial KNN graph. and estimated a GFTscore per gene on the Fourier coefficient for low-frequency signals. We installed *SpaGFT* (v0.1.1.4) and ran it by following the tutorial: https://spagft.readthedocs.io/en/latest. We computed auPRC for this method using GFTscore and selected significant SVGs using q-value < 0.05.

#### SpaGCN

*SpaGCN* is a graph convolutional network (GCN)-based approach that integrates gene expression, spatial location, and histology to identify SVGs^29^. It first identifies spatial domains through clustering and then detects SVGs that are enriched in each domain. We installed *SpaGCN* (v1.2.5) and ran the method by following the tutorial: https://github.com/jianhuupenn/SpaGCN/blob/master/tutorial/tutorial.ipynb. We used the adjusted p-values to rank genes for computing auPRC and select significant SVGs (<0.05).

#### scGCO

*scGCO* (single-cell graph cuts optimization) utilizes a hidden Markov random field (HMRF) with graph cuts to identify SVGs^27^. We installed *scGCO* (v1.1.0) and executed the method by following the tutorial: https://github.com/WangPeng-Lab/scGCO/blob/master/code/Tutorial/scGCO_tutorial.ipynb. To compute auPRC, we used FDR to rank the genes. To select significant SVGs, we used 0.05 of FDR as threshold.

#### Sepal

*Sepal* assesses the degree of randomness exhibited by the expression profile of each gene through a diffusion process and ranks the genes accordingly^31^. We computed the *Sepal* score using Squidpy (v1.2.3) by following the tutorial: https://squidpy.readthedocs.io/en/stable/auto_examples/graph/compute_sepal.html. We used the sepal score to rank the genes to calculate auPRC.

#### SpatialDE

*SpatialDE* is one of the pioneer methods for identifying SVGs^32^. It models the normalized spatial gene expression using the Gaussian process regression and estimates the significance by comparing the models with and without spatial covariance. We installed *SpatialDE*∼(v1.1.3) with pip and processed the data with the functions NaiveDE.stabilize and NaiveDE.regress_out. We ran the function SpatialDE.run to obtain results and used the fraction of spatial variance (FSV) to compute auPRC. To select significant SVGs, we used the adjusted p-values (< 0.05).

#### SpatialDE2

*SpatialDE2* is a flexible framework for modeling spatial transcriptomics data that refines *SpatialDE* by providing technical innovations and computational speedups^23^. We obtained the source code from https://github.com/PMBio/SpatialDE and estimated spatial variance using the function SpatialDE.fit. Similar to *SpatialDE*, we ranked the genes by FSV to compute auPRC.

#### SPARK

*SPARK* extended the computation framework proposed in SpatialDE by directly modeling the raw count data using a generalized linear spatial model (GLSM) based on Poisson distribution^33^. We obtained *SPARK* (v1.1.1) from https://github.com/xzhoulab/SPARK and ran the method by following the tutorial https://xzhoulab.github.io/SPARK/02_SPARK_Example. We used the adjusted p-values to compute auPRC and select significant SVGs (< 0.05).

#### SPARK-X

*SPARK-X* is a non-parametric covariance test method based on multiple spatial kernels for modeling sparse count data from spatial transcriptomic experiments^37^. We ran *SPARK-X* (v1.1.1) by following the tutorial: https://xzhoulab.github.io/SPARK/02_SPARK_Example. We used the adjusted p-values to compute auPRC and select significant SVGs (< 0.05).

#### BOOST-GP

*BOOST-GP* is a Bayesian hierarchical model to analyze spatial transcriptomics data based on zero-inflated negative binomial distribution and Gaussian process^35^. We downloaded the source codes of *BOOST-SP* from https://github.com/Minzhe/BOOST-GP and ran the function boost.gp by setting the parameters iter to 100 and burn to 50. We used p-values to compute auPRC and select significant SVGs using 0.05 as the threshold.

#### GPcounts

*GPcounts* implemented Gaussian process regression for modeling counts data using a negative binomial likelihood function^36^. We obtained the source codes of *GPcounts* from https://github.com/ManchesterBioinference/GPcounts and followed the tutorial https://github.com/ManchesterBioinference/GPcounts/blob/master/demo_notebooks/GPcounts_spatial.ipynb. To compute auPRC, we ranked the genes by the log-likelihood ratio (LLR), representing the ratio between the dynamic and constant (null) models. Significant SVGs were selected based on the q-values with 0.05 as the threshold. We noted that *GPcounts* sometimes failed to generate results for certain genes, especially when applied to real-world datasets. In this case, we set the LLR as 0 and the q-value as 1.

#### nnSVG

*nnSVG* is a method built on nearest-neighbor Gaussian processes to identify SVGs^38^. We installed the package (v1.2.0) from Bioconductor and ran the method by following the tutorial https://bioconductor.org/packages/release/bioc/vignettes/nnSVG/inst/doc/nnSVG.html. We used the fraction of spatial variance estimated by the method to compute auPRC. Significant SVGs were selected based on adjusted p-values using 0.05 as the threshold.

#### SOMDE

*SOMDE* uses a self-organizing map (SOM) to cluster neighboring locations into nodes and then uses a Gaussian process to fit the node-level spatial gene expression to identify SVGs^39^. We installed *SOMDE* (v0.1.7) with pip and followed the tutorial https://github.com/WhirlFirst/somde/blob/master/slide_seq0819_11_SOM.ipynb to run the method. We ranked the genes by FSV to compute auPRC and selected significant SVGs based on the q-values using 0.05 as the threshold.

### Benchmarking prediction performance, sensitivity, and specificity

We applied each method on the simulated datasets to identify SVGs. For comparison, we computed the auPRC using the function pr.curve from the R package PRROC^52^ by ranking the prediction for each method accordingly (**see Table 1**). We calculated the true positive rate (sensitivity) and true negative rate (specificity) at the false discovery rate of 0.05 as follows:

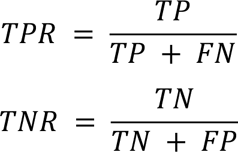

where *TP* denotes the number of true positives, *FN* denotes the number of false negatives, *TN* denotes the number of true negatives and *FP* denotes the number of false positives. For SpatialDE2 and Sepal, we selected the top n genes (n = the number of true positives) as significant SVGs to sensitivity and specificity.

### Benchmarking scalability with the number of spatial spots

We used the Snakemake^53^ workflow (v7.25.2) management system to evaluate the scalability of each method with the number of spatial spots. For this, we generated simulation datasets with 100 genes and various numbers of spots from 100 to 40000 (see above). Next, we ran each method on a dedicated HPC node with AMD EPYC 7H12 64-Core Processor using the same computational resource (1TB memory, 120 hours, and 10 CPUs) defined by the Snakemake pipeline. For methods (i.e., *GPcounts* and *SpatialDE2*) that require a graphics processing unit (GPU) for running, we used an A100 with 40GB of memory. We measured the memory usage and running time using the benchmark directive provided by the Snakemake tool (--benchmark). Notably, we could not run *SPARK* for datasets with 40000 spots because of memory issues. Moreover, *BOOST-GP* did not generate results for datasets with 20000 and 40000 spots within 120 hours.

### Benchmarking impact of identified SVGs on spatial domain detection analysis

We utilized the human DLPFC datasets to evaluate the impact of **identified** SVGs on spatial domain detection. We ran the methods and determined the SVGs based on an FDR 0.05. Since *SpatialDE2* and *Sepal* did not provide statistical significance, we selected the top 2000 genes based on the FSV and Sepal scores, respectively. Because *BOOST-GP* failed to produce any results after 120 hours of running, we excluded it from this evaluation. In addition, we also identified the highly variable genes using the function scanpy.pp.highly_variable_genes and used the top 2000 as our baseline for comparison. We next used these genes to perform dimension reduction using the function scanpy.tl.pca and generated a k-nearest-neighbor graph with scanpy.pp.neighbors. The clustering was conducted using the function scanpy.tl.leiden (resolution = 1). We next compared the obtained clusters (denoted by *X*) with the annotated layers (denoted by *Y*). We assessed the clustering quality with Adjusted Rand Index (ARI):

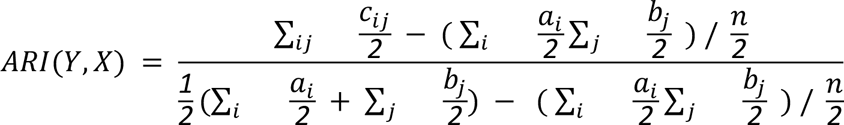

where *c*_*ij*_ denotes the number of common spots for each obtained cluster *i* and ground truth *j*, 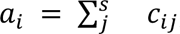, and 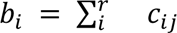. We also calculated the Normalized Mutual Information (NMI) for comparison as follows:

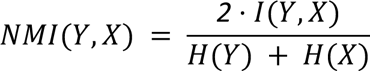

where *H* represents entropy of the partition and *I* represents the mutual information between clusters and the true labels. Both ARI and NMI have values from 0 to 1, with 1 indicating that the two partitions are the same and 0 indicating that the two are independent.

### Benchmarking the methods for spatial ATAC-seq data

We downloaded spatial ATAC-seq data of mouse embryos at stages E12.5, E13.5, and E15.5 from GEO with accession number GSE214991. We first identified open chromatin regions by peak calling on all the spots using MACS2^54^ (--nomodel --nolambda --shift -75 --extsize 150) and obtained 34460 (E12.5), 31099 (E13.5), and 69896 (E15.5) peaks for each sample, respectively. We next built a cell-by-peak count matrix using the fragments and peaks as input based on the function FeatureMatrix from the Signac^55^ package. We only retained the spots that were located on the tissue.

We ran each method on the cell-by-peak matrix of spatial ATAC-seq data from mouse embryos to detect spatially variable peaks. For those methods that require normalized data as input, we used TF-IDF (Term Frequency - Inverse Document Frequency) for normalization. Of note, *BOOST-GP* and *GPcounts* failed to produce results after 120 hours, and we could not obtain results from *SPARK* due to memory issues. Therefore, these three methods were excluded from this evaluation. We selected the significant variable peaks using an FDR of 0.05 for the rest of the methods. For *SpatialDE2* and *Sepal*, we opted for the top 20000 peaks since they did not provide statistical significance. We next used the spatially variable peaks to cluster the spots. Because the true clusters were unavailable, we evaluated the clustering performance by following ref.^18^ based on the local inverse Simpson’s index (LISI) and the spatial chaos score (CHAOS). The LISI score was calculated as follows:

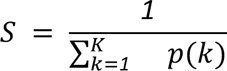

where *p(k)* is the probability that the cluster label *k* is in the local neighborhood, and *K* is the total number of clusters. A lower LISI score indicates more homogeneous neighborhood clusters of the spots. The CHAOS score was calculated as follows:

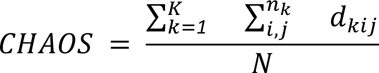

where *d_kij_* is the Euclidean distance between the spots *i* and *j* in the cluster *k* and *N* is the total number of spots. A lower CHAOS indicates better spatial continuity.

## Data Availability

All the datasets used in this study are publicly available. The human DLPFC data were downloaded from http://research.libd.org/spatialLIBD. The breast tumor data were downloaded from https://github.com/almaan/her2st. Spatial-ATAC-seq data were obtained from the Gene Expression Omnibus (GEO) with the accession number GSE214991.

## Code Availability

The code for running the benchmarked methods is available on GitHub: https://github.com/pinellolab/SVG_Benchmarking. The code for generating the simulation datasets is available on Github: https://github.com/pinellolab/simstpy.

## Acknowledgment

The authors would like to thank the members of Pinello Lab for the discussion. L.P. is partially supported by the National Human Genome Research Institute (NHGRI) Genomic Innovator Award (R35HG010717).

## Author contributions statement

Z.L. and L.P. conceived the study. Z.L. and Z.P. conducted the experiments and analyzed the results. D.S. and G.Y. supported the simulation results using scDesign3. Z.L. and Z.P. wrote the manuscript, revised by L.P. and J.J.L. All authors reviewed the manuscript.

## Notes

### Competing Interest Statement

The authors have declared no competing interest.

