## Supplemental Figures for "Benchmarking computational methods to identify spatially variable genes and peaks"

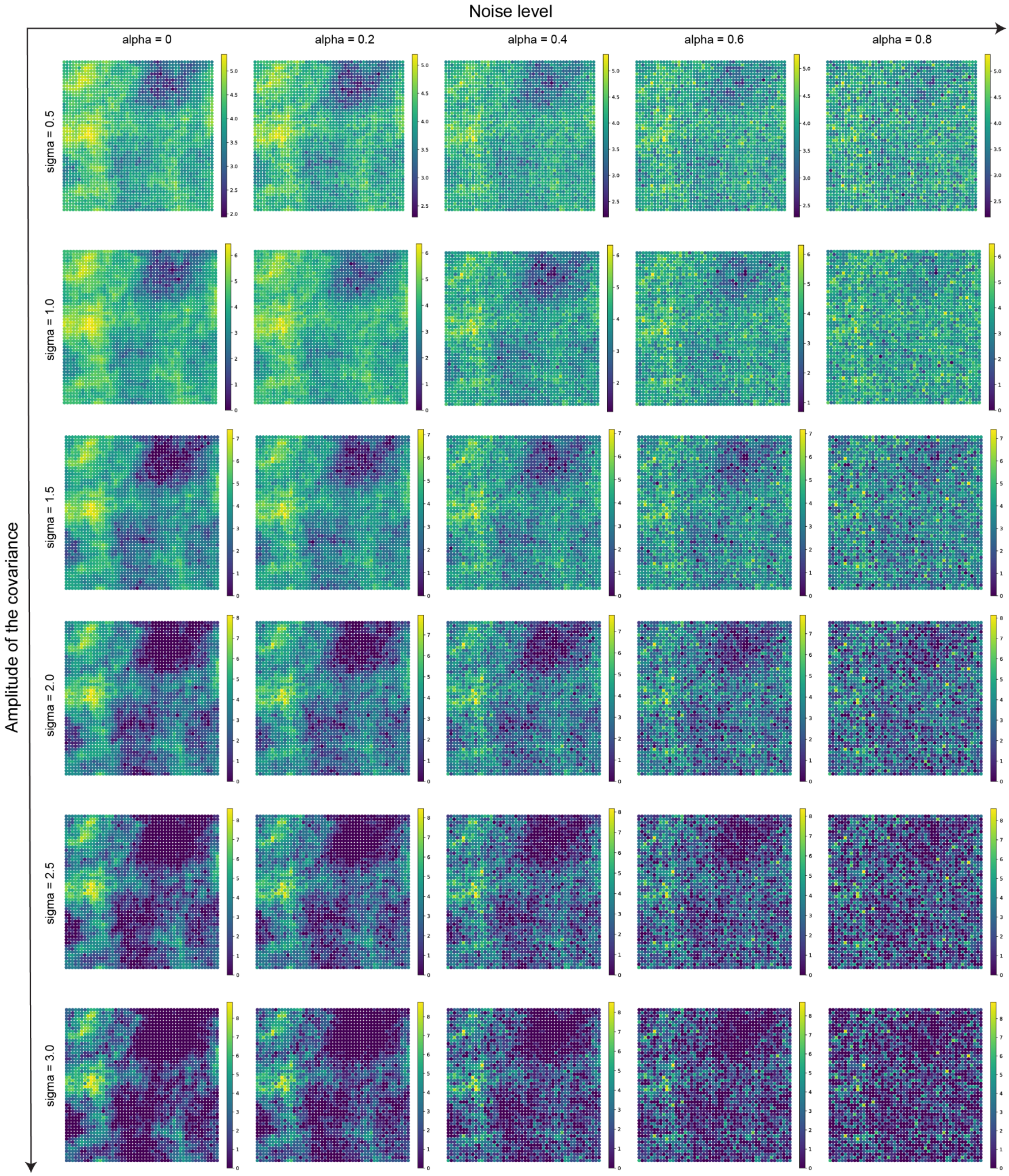


**Supplementary Fig. 1 Visualization of covariance-based simulation.** Each plot represents a simulated spatially variable gene with different noise levels (denoted by alpha) and amplitude of the covariance matrix (denoted by sigma). Colors refer to normalized expression profiles. X-axis and y-axis represent spatial coordinates.

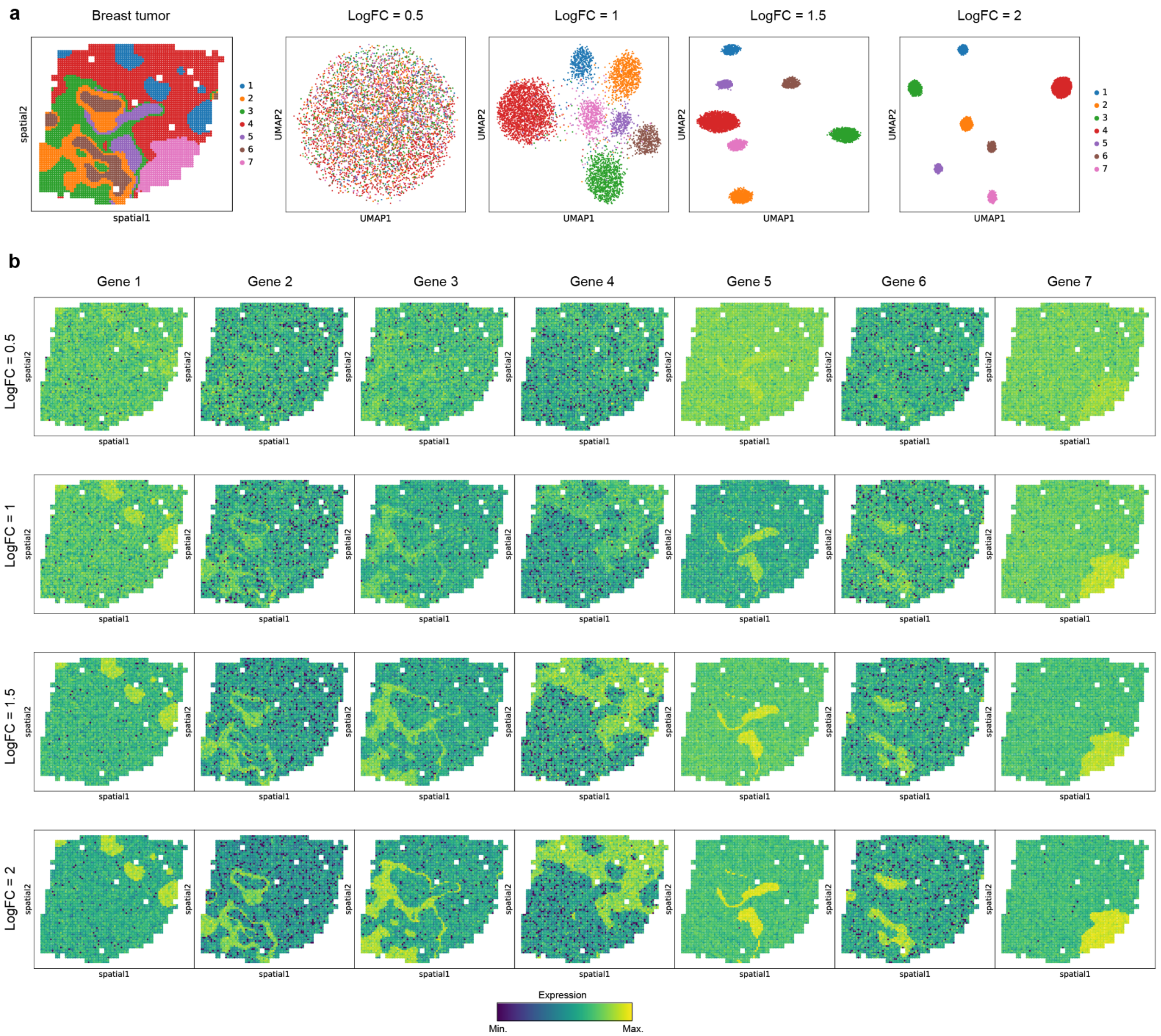


**Supplementary Fig. 2 Visualization of clustering-based simulation. a,** Simulated data based on breast cancer. Left: Visualization of annotated tissue structures in spatial space. Colors refer to different clusters. Right: UMAP embedding of all spots generated using only spatially variable genes with different log fold changes from 0.5 to 2. **b,** Visualization of the spatially variable gene for each cluster. Each column represents a gene, and each row represents a specific log fold change.

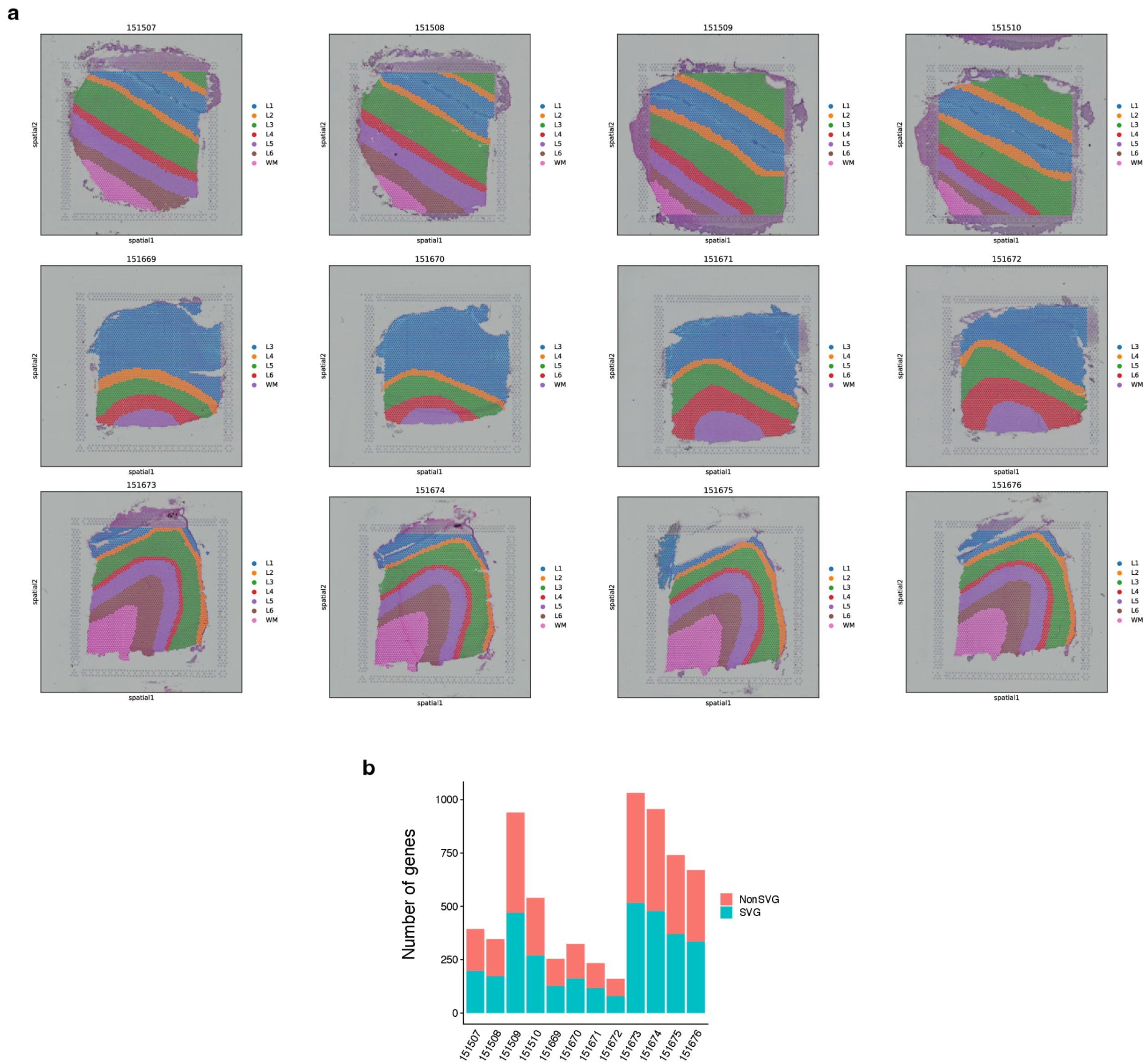


**Supplementary Fig. 3 Visualization of the shuffling-based simulation datasets from human DLPFC. a,** Visualization of the 12 DLPFC datasets. ID for each dataset is shown on the top. The colors refer to cortex layers, which were manually annotated. **b,** The number of genes per data is shown as a stacked bar plot. Colors refer to true positives and true negatives.

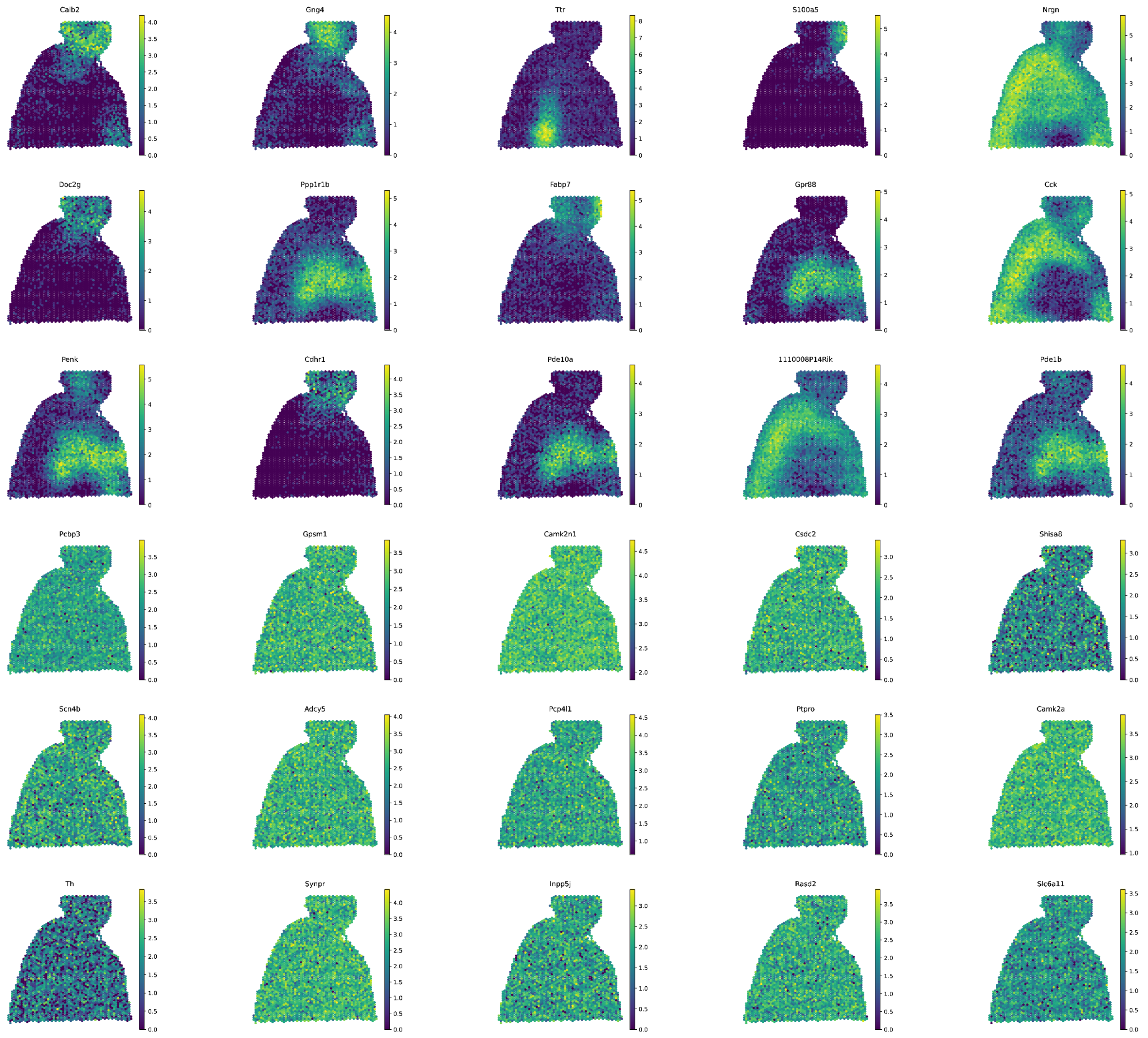


**Supplementary Fig. 4 Visualization of scDesign3 simulated data.** Simulated data was generated by using the scDesign3 pipeline. The top 3 rows represent true spatially variable genes, and the bottom three rows represent non-spatially variable genes.

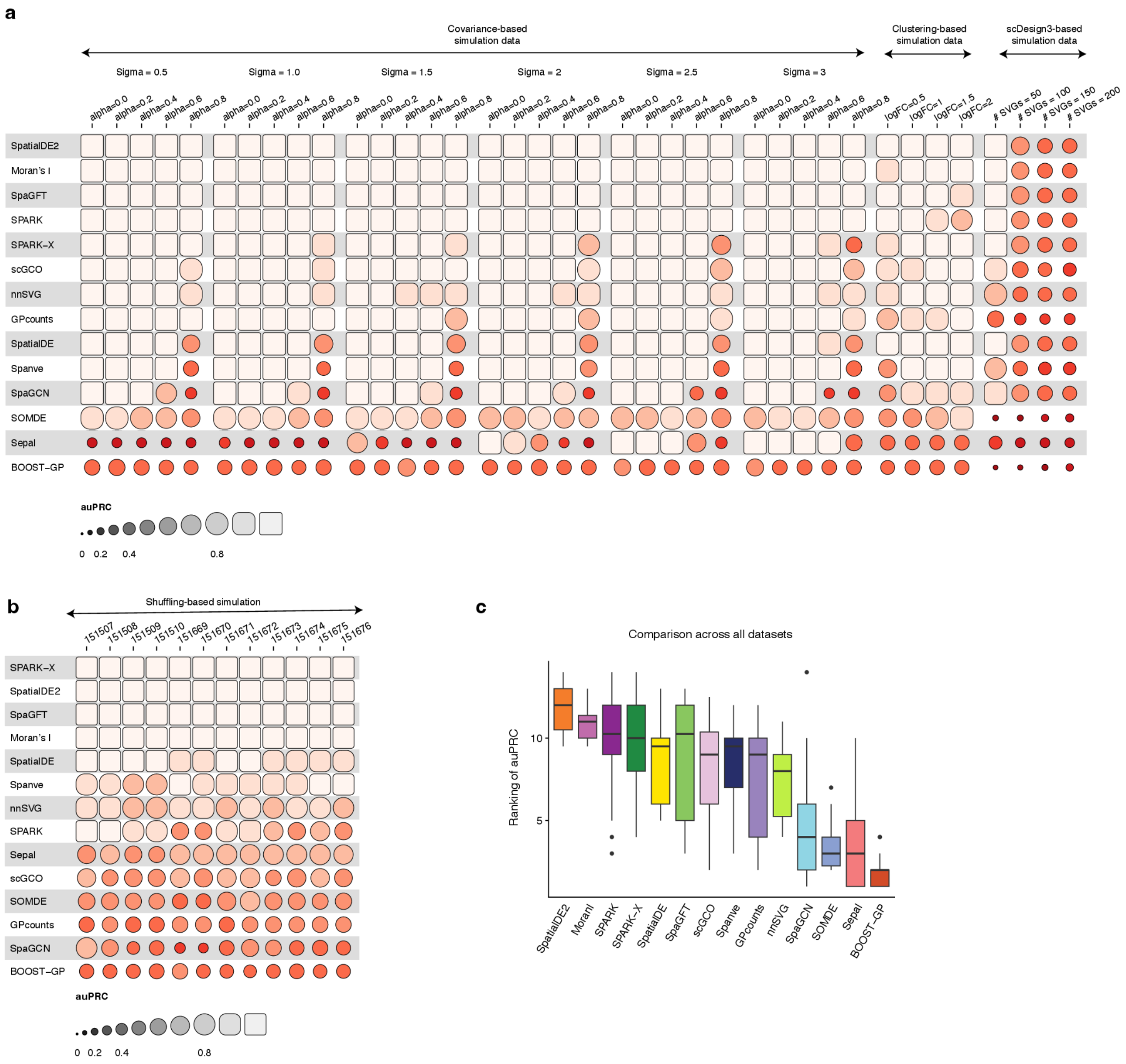


**Supplementary Fig. 5 Evaluated prediction accuracy of the methods. a,** Comparison of the methods as evaluated by auPRC using simulation datasets. Each row represents a method, and each column represents a dataset. Methods are ranked based on the mean auPRC across the datasets. **b,** Same as **a** for the shuffling-based simulation data. **c,** Boxplot showing the overall ranking of the methods across all datasets (i.e., covariance-based, clustering-based, scDesign3-based, and shuffling-based simulation datasets). Related to **Fig. 2a**.

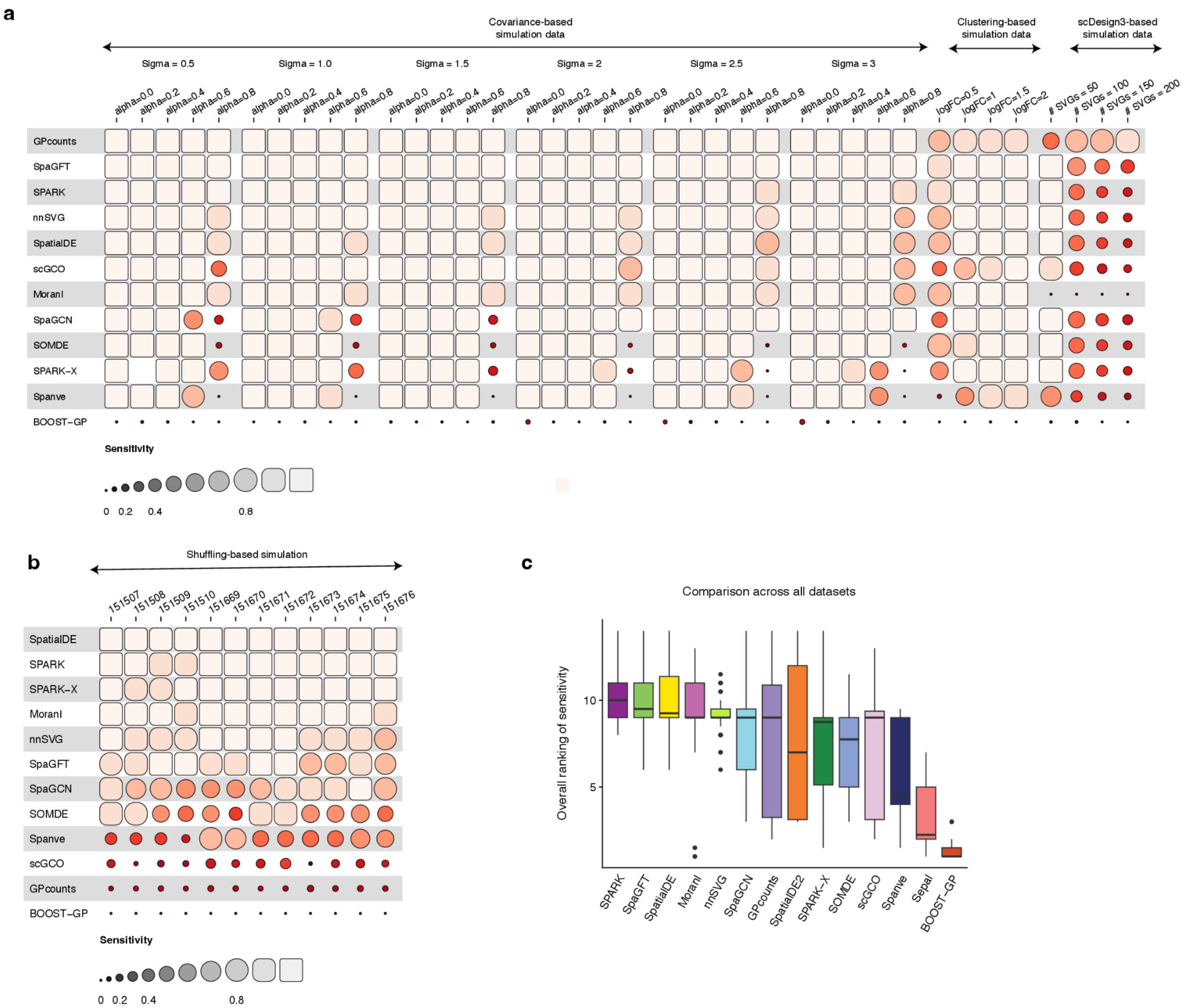


**Supplementary Fig. 6 Comparison of the sensitivity of the methods. a,** Comparison of sensitivity between the methods using simulation datasets. Each row represents a method, and each column represents a dataset. Methods are ranked based on the mean sensitivity across the datasets. **b,** Same as **a** for the shuffling-based simulation data. **c,** Boxplot showing the overall ranking of the methods across all datasets (i.e., covariance-based, clustering-based, scDesign3-based, and shuffling-based simulation datasets). Related to **Fig. 2b**.

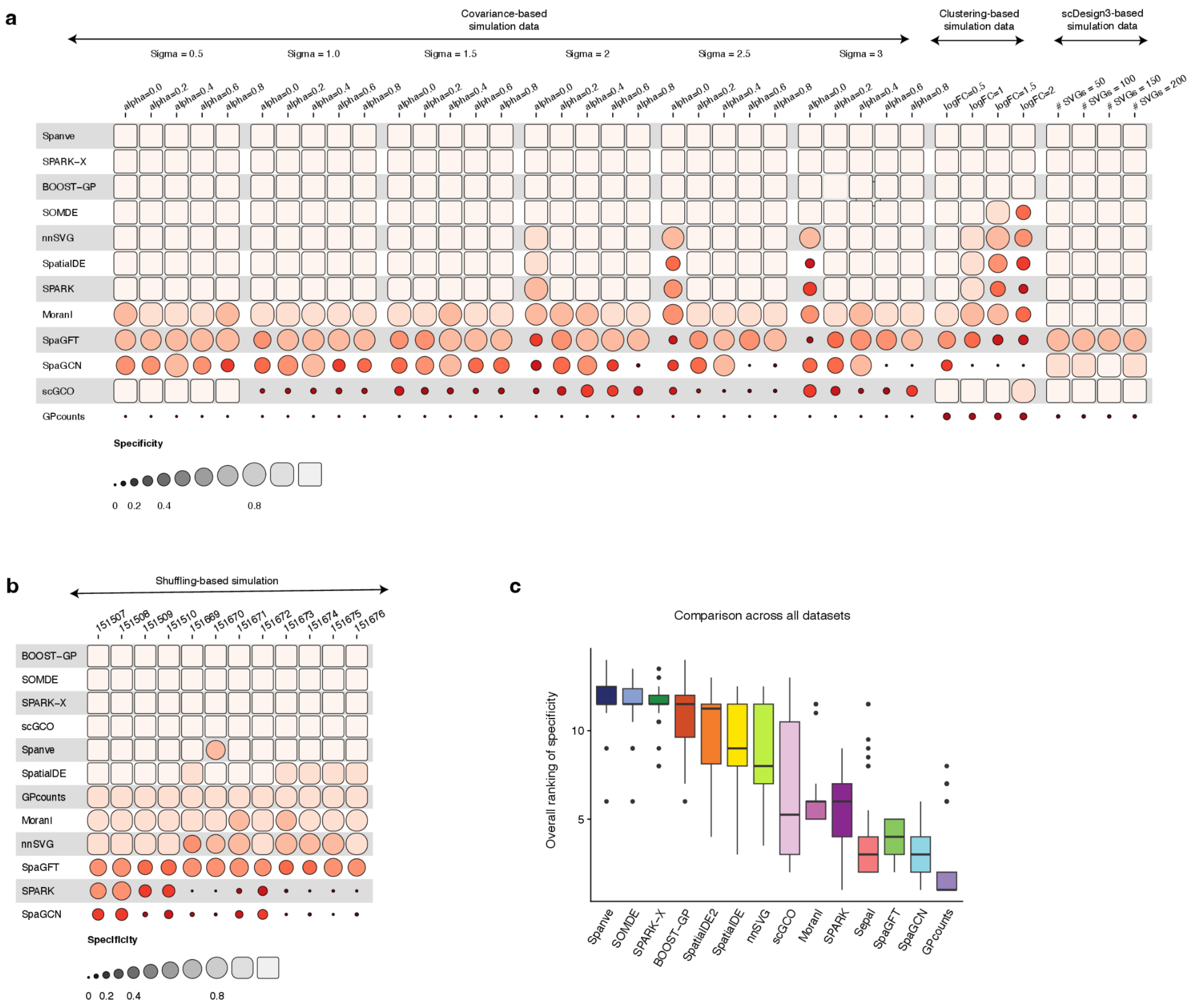


**Supplementary Fig. 7 Comparison of the specificity of the methods. a,** Comparison of specificity between the methods using simulation datasets. Each row represents a method, and each column represents a dataset. Methods are ranked based on the mean specificity across the datasets. **b,** Same as **a** for the shuffling-based simulation data. **c,** Boxplot showing the overall ranking of the methods across all datasets (i.e., covariance-based, clustering-based, scDesign3-based, and shuffling-based simulation datasets). Related to **Fig. 2b**.

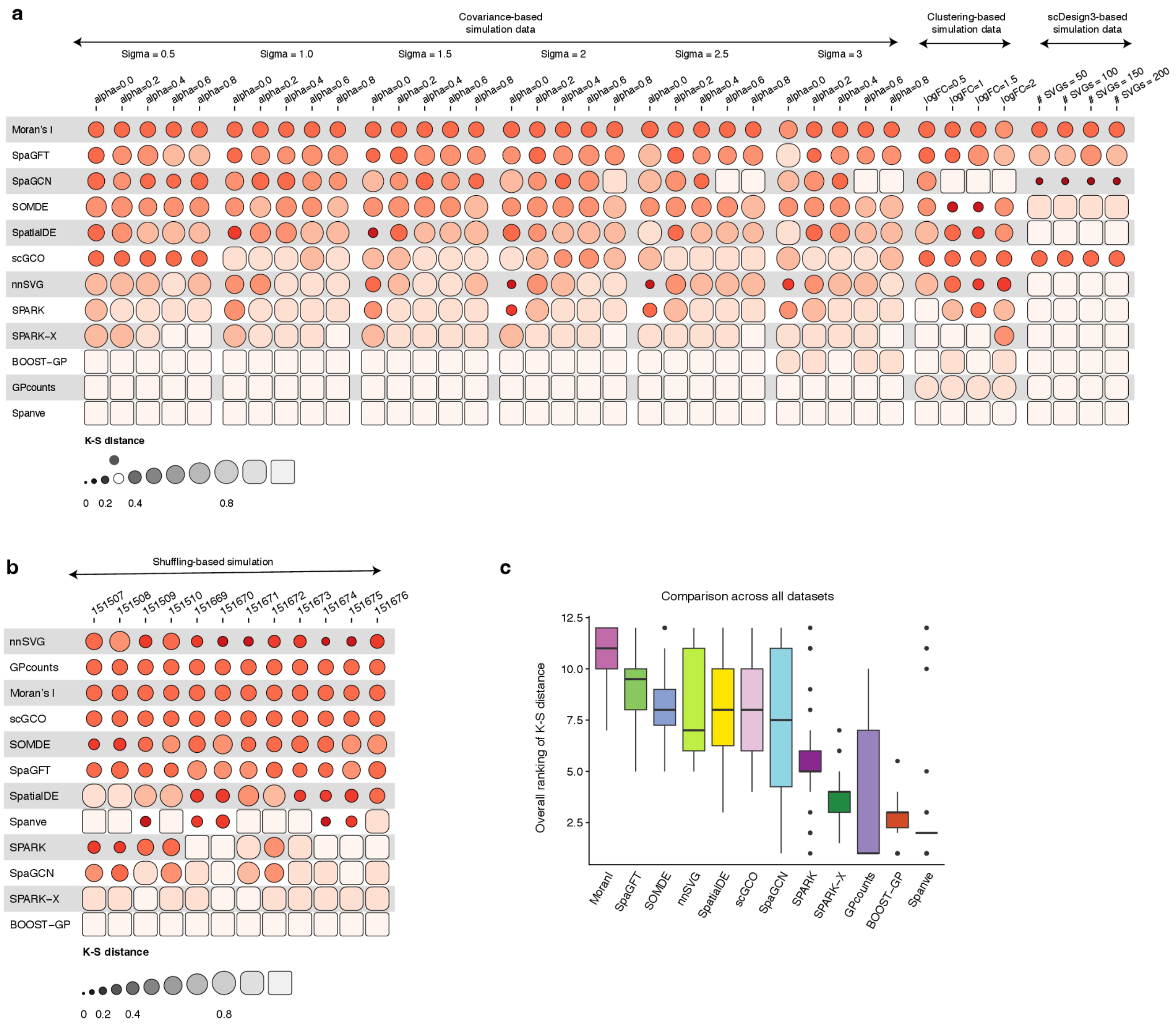


**Supplementary Fig. 8 Comparison of the statistical calibration of the methods. a,** Comparison of p-value calibration as measured by K-S distance between the methods using simulation datasets. Each row represents a method, and each column represents a dataset. Methods are ranked based on the mean K-S distance across the datasets. **b,** Same as **a** for human DLPFC dataset. **c,** Boxplot showing the overall ranking of the methods across all datasets (i.e., covariance-based, clustering-based, scDesign3-based, and shuffling-based simulation datasets). Related to **Fig. 2c**.

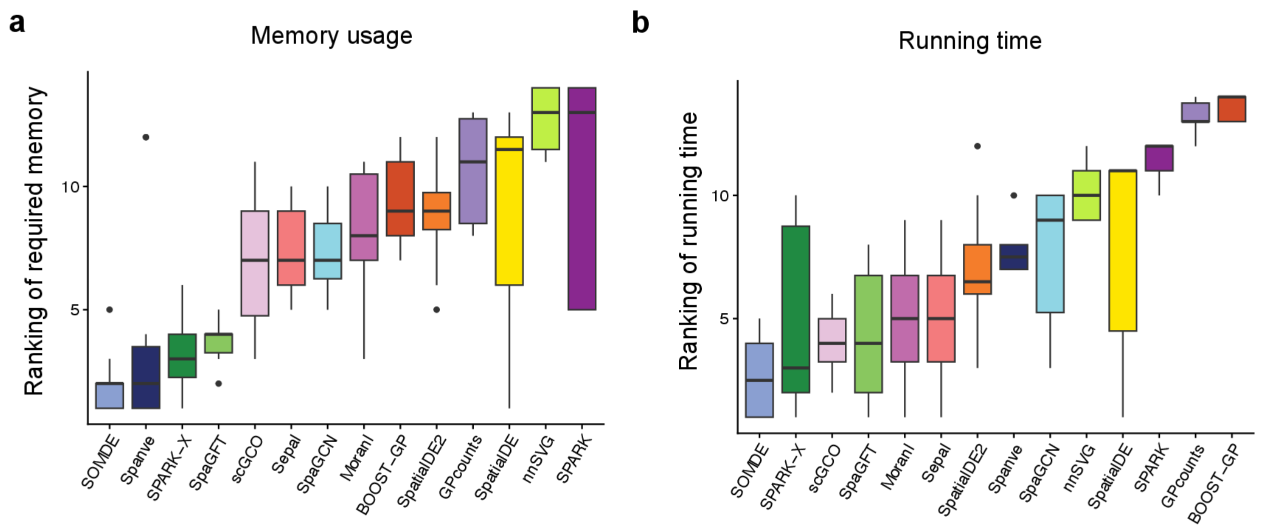


**Supplementary Fig. 9 Comparison of memory and running time between methods. a,** Boxplot showing the ranking of required memory by each method across the benchmarked datasets. Lower rank represents less requirement of memory. **b,** Same as **a** for running time.

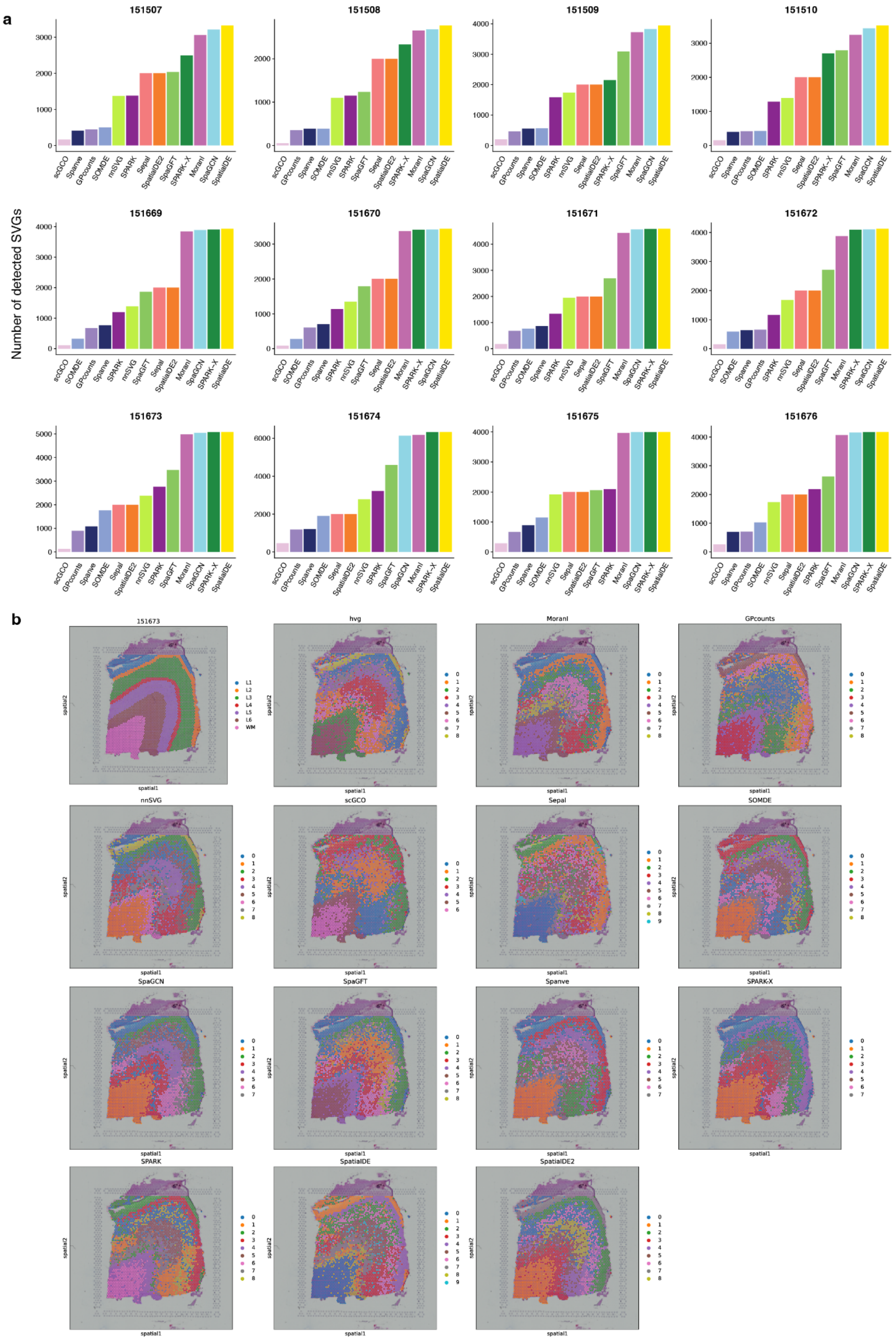
a
151507
151508
151509
151510
151669
151670
151671
151672
151673
151674
151675
151676
Number of detected SVGs
scGCO
Spanve
GPcounts
SOMDE
nnSVG
SPARK
Sepal
SpatialDE2
SpaGFT
SPARK-X
MoranI
SpaGCN
SpatialDE
b
151673
hvg
MoranI
GPcounts
nnSVG
scGCO
Sepal
SOMDE
SpaGCN
SpaGFT
Spanve
SPARK-X
SPARK
SpatialDE
SpatialDE2
spatial1
spatial2
L1
L2
L3
L4
L5
L6
WM

**Supplementary Fig. 10 Impact of SVGs on clustering analysis. a,** Bar plots showing the number of detected SVGs for each method on different human DLPFC datasets. **b,** Visualization of the clustering results for dataset 151673.

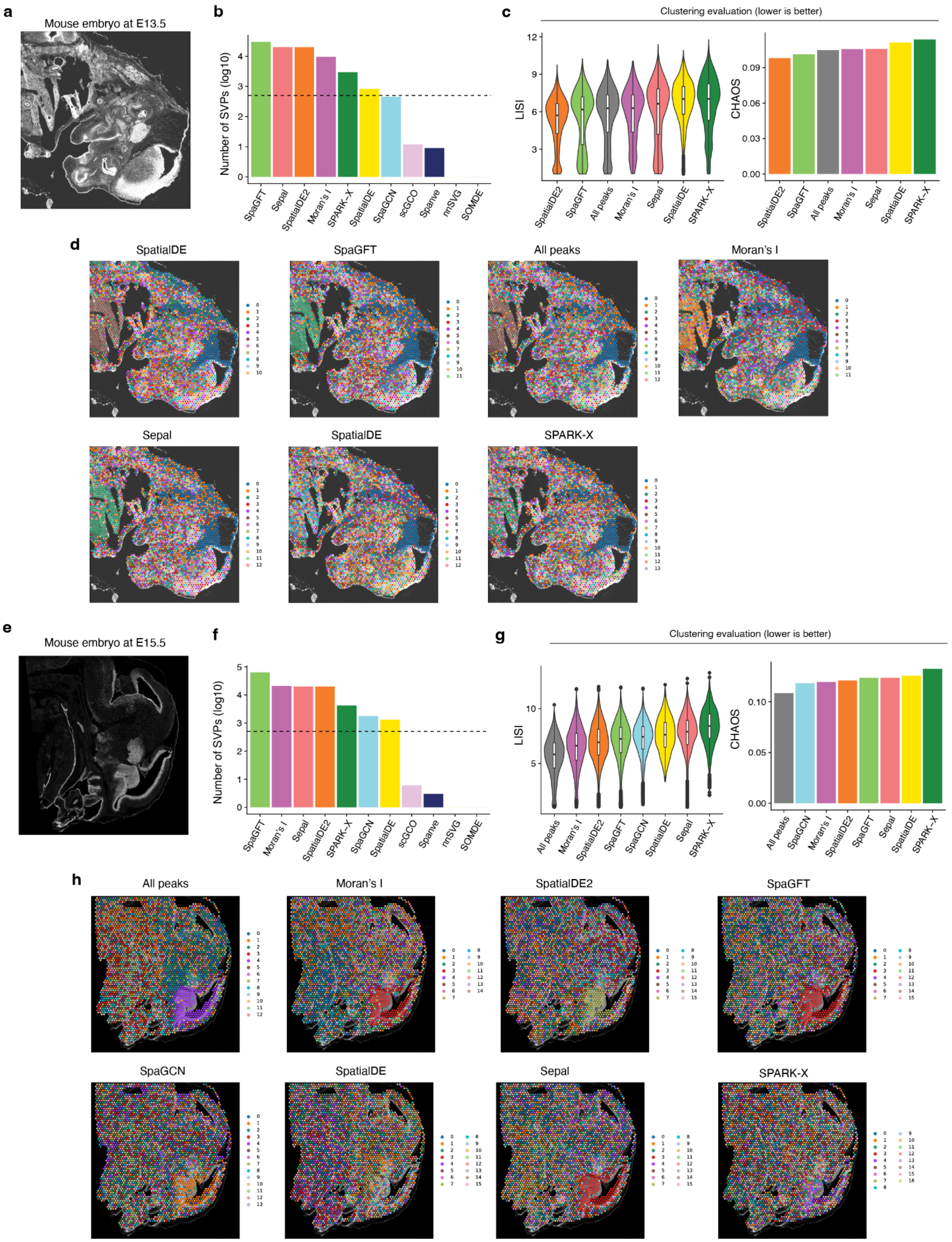
a
Mouse embryo at E13.5
b
Number of SVPs (log10)
SpaGFT
Sepal
SpatialDE2
Moran's I
SPARK-X
SpatialDE
SpaGCN
scGCO
Spanve
nnSVG
SOMDE
c
Clustering evaluation (lower is better)
LISI
CHAOS
All peaks
d
SpatialDE
SpaGFT
All peaks
Moran's I
Sepal
SpatialDE
SPARK-X
e
Mouse embryo at E15.5
f
Number of SVPs (log10)
g
Clustering evaluation (lower is better)
LISI
CHAOS
h
All peaks
Moran's I
SpatialDE2
SpaGFT
SpaGCN
SpatialDE
Sepal
SPARK-X

**Supplementary Fig. 11 Benchmarking the methods using spatial ATAC-seq. a,** Image of a mouse embryo at days of E13.5. **b,** Number of detected spatially variable peaks by each method. The dashed line represents the cutoff of the number of peaks (n=500) used for downstream clustering analysis. Methods are sorted by the number of detected peaks. **c,** Left: violin plot showing the LISI scores. Methods are sorted by the median values. Right: The bar plot shows the CHAOS score. For both metrics, a lower value represents a better performance. **d,** Visualization of obtained clusters using spatially variable peaks identified by different methods. **e,** Same as **a** mouse embryo at days of E15.5. **f,** Same as **b** for mouse embryo at days of E15.5. **g,** Same as **c** for mouse embryos at days of E15.5. **h,** Same as **d** for mouse embryo at days of E15.5.
